# Cell-Specific Gene Networks and Drivers in Rheumatoid Arthritis Synovial Tissues

**DOI:** 10.1101/2023.12.28.573505

**Authors:** Aurelien Pelissier, Teresina Laragione, Percio S. Gulko, María Rodríguez Martínez

**Affiliations:** IBM Research Europe, 8803 Rüschlikon, Switzerland; Department of Biosystems Science and Engineering, ETH Zurich, 4058 Basel, Switzerland; Currently at Institute of Computational Life Sciences, ZHAW, 8400 Winterthur, Switzerland; Division of Rheumatology, Icahn School of Medicine at Mount Sinai, 10029 New York, United States; Currently at Yale School of Medicine, 06510 New Haven, United States

**Author notes:** P. S. Gulko contributed equally with M. R. Martinez.

## Abstract

Rheumatoid arthritis (RA) is a common autoimmune and inflammatory disease characterized by inflammation and hyperplasia of the synovial tissues. RA pathogenesis involves multiple cell types, genes, transcription factors (TFs) and networks. Yet, little is known about the TFs, and key drivers and networks regulating cell function and disease at the synovial tissue level, which is the site of disease. In the present study, we used available RNA-seq databases generated from synovial tissues and developed a novel approach to elucidate cell type-specific regulatory networks on synovial tissue genes in RA. We leverage established computational methodologies to infer sample-specific gene regulatory networks and applied statistical methods to compare network properties across phenotypic groups (RA versus osteoarthritis). We developed computational approaches to rank TFs based on their contribution to the observed phenotypic differences between RA and controls across different cell types. We identified 18,16,19,11 key regulators of fibroblast-like synoviocyte (FLS), T cells, B cells, and monocyte signatures and networks, respectively, in RA synovial tissues. Interestingly, FLS and B cells were driven by multiple independent co-regulatory TF clusters that included MITF, HLX, BACH1 (FLS) and KLF13, FOSB, FOSL1 (synovial B cells). However, monocytes were collectively governed by a single cluster of TF drivers, responsible for the main phenotypic differences between RA and controls, which included RFX5, IRF9, CREB5. Among several cell subset and pathway changes, we also detected reduced presence of NKT cell and eosinophils in RA synovial tissues. Overall, our novel approach identified new and previously unsuspected KDG, TF and networks and should help better understanding individual cell regulation and co-regulatory networks in RA pathogenesis, as well as potentially generate new targets for treatment.

## Introduction

Rheumatoid arthritis (RA) is a systemic autoimmune disorder characterized by synovial inflammation and hyperplasia that may lead to joint destruction [1, 2]. With a prevalence estimated between 0.5 and 1% [3], it is one of the most common chronic inflammatory diseases. The risk of developing RA peaks around age 50 [3], and similarly to most autoimmune diseases, females are affected 2 to 3 times more than males [4, 3]. The development of biologic disease-modifying anti-rheumatic drugs (bDMARDs) and JAK inhibitors targeting various inflammatory pathways has significantly improved disease control and outcomes [5, 6], yet a considerable number of RA patients still have an inadequate response to therapy [7]. As the development and progression of RA involve dynamic interactions between multiple genetic and environmental factors, understanding the heterogeneous pathophysiological processes remains a major challenge [8]

Genome-Wide Association Studies (GWAS) have significantly improved the understanding of the disease’s genetic underpinnings and identified multiple genetic loci associated with susceptibility [9, 10]. However, these loci only explain a fraction of the overall heritability and phenotypic variance of RA [11]. While new whole-genome and whole-exome sequencing studies are likely to identify additional rare variants previously undetectable with GWAS techniques, our ability to translate these results into disease understanding and novel therapeutics remains limited. At the transcriptomic level, Differential Gene Expression (DGE) studies have compared gene expression profiles between RA patients and healthy controls and identified numerous pathways implicated in inflammation, antigen presentation, hypoxia, and apoptosis during RA [12, 13]. These data have not only deepened the comprehension of RA’s pathogenesis but have also offered promising targets for therapeutic intervention. However, although DGE studies are effective for discovery, the large number of detected genes often obscures the identification of key regulatory or therapeutically actionable genes. Moreover, DGE studies often highlight pathways that are already well-characterized, underscoring the need for new approaches to integrate established knowledge and data-driven computational approaches.

Further complicating the analysis, most studies still rely on bulk data, wherein the cellular composition of tissues significantly confounds the molecular findings [14, 15]. In other words, the observed differential expression between RA and controls may primarily arise from disparities in cell composition rather than differences in cellular gene expression. For instance, RA synovial tissues exhibit significantly more leukocytes than control groups [15, 16], and a meta-analysis has revealed variable DEGs across datasets, with approximately half of the DEGs up-regulated in synovial tissues being down-regulated in blood samples [12]. Recent single-cell RNA sequencing (scRNA-Seq) sequencing studies are offering valuable insights into disease traits at the single-cell level [15, 16], however, the exploitation of these data is still challenged by limited patient numbers, batch effects, and sparse data [17]. Notably, the inference of gene regulatory networks (GRNs) from scRNA-Seq has proven to be particularly challenging, in part because of the difficulty of identifying cell type-specific regulatory interactions from heterogeneous samples [18, 19, 20].

Addressing the cell composition challenge requires the development of novel approaches for identifying transcription factors (TFs) and their associated regulatory signatures in a cell type-specific manner. TFs play a pivotal role in regulating gene expression in a tissue-specific manner [21], and with an estimated count of 1,000 TFs in humans [22], identifying those that govern the phenotypic traits of RA synovial cells may offer opportunities for discovering novel therapeutic targets.

In this study, we provide a comprehensive analysis of gene regulation of RA in synovial tissues. Unlike previous studies in RA, which relied on the inference of cohort-averaged GRNs [23, 24, 25], our approach leveraged bulk gene expression data [26, 15] and enabled us enables us to gain new insights into sample-specific regulatory mechanisms and to identify TFs driving gene expression in RA-associated cell types. Namely, we identify key RA regulators in a cell type-specific manner, such as IRF8 in monocytes, STAT5B in B cells, ELF4 in T cells, and MITF in fibroblast-like synoviocytes (FLS). We next construct a co-regulation network in each cell type by evaluating the correlation among the target genes that are shared by the identified TFs. In FLS and B cells, we observe a common regulatory pattern characterized by a strong correlation among all TFs, while distinct co-regulation clusters emerge independently in the other cell types. Our computational modeling and findings indicate new targets for cell-specific treatment strategies in RA and provide novel insights into the cell-specific regulation of RA pathogenesis.

## 1 Results

### 1.1 Heterogeneous cellular composition accounts for most of the gene expression variability across RA biopsies

We exploited analyzed public RNA-Seq data of synovial biopsies from 28 healthy control samples, 152 individuals with RA, and 22 patients with osteoarthritis (OA) [26]. Each sample contained the gene expression profile of 25,000 genes. As healthy biopsies are rare and difficult to obtain [27], we compared RA synovial biopsies to both OA and healthy samples

We first used UMAP [28] to visualize the samples in two dimensions, and observed significant distributional differences between the gene expression profiles of RA biopsies and controls, the latter comprising OA and healthy samples (Figure 1A). A DEG analysis revealed that more than half of the genes (∼15k) were significantly differentially expressed (p < 0.05; Student’s t-test with Benjamini-Hochberg correction [29]). This uncommonly high number of DEGs, which is typically on the order of a few hundred in studies of blood samples from patients with other diseases, such as coronary artery disease [30], obesity [31, 32], diabetes [33, 34] or kidney [35], led us to hypothesize that the observed variability might arise from cellular heterogeneity across synovial biopsies, rather than from intra-cellular gene expression differences between the two tested groups. To test this hypothesis, we used xCell [36] to estimate the relative proportions of the various cell populations present in our samples. xCell is a signature-based method that employs single-sample gene set enrichment analysis to compute an enrichment score per sample. This score is indicative of the relative proportion of each cell type within a sample (Methods Section 2.2). xCell detected significantly increased enrichment scores for several cell populations in both RA vs normal and RA vs OA tissue comparisons (Figure 1B; *p* < 0.05, Student’s *t*-test). Interestingly, biopsies from early and established RA had similar gene expression signatures and cellularity, suggesting that similar cell types and processes regulate disease through the different stages of progression (Supplementary Section A.1, Supplementary Figure S1).

**Figure 1:**
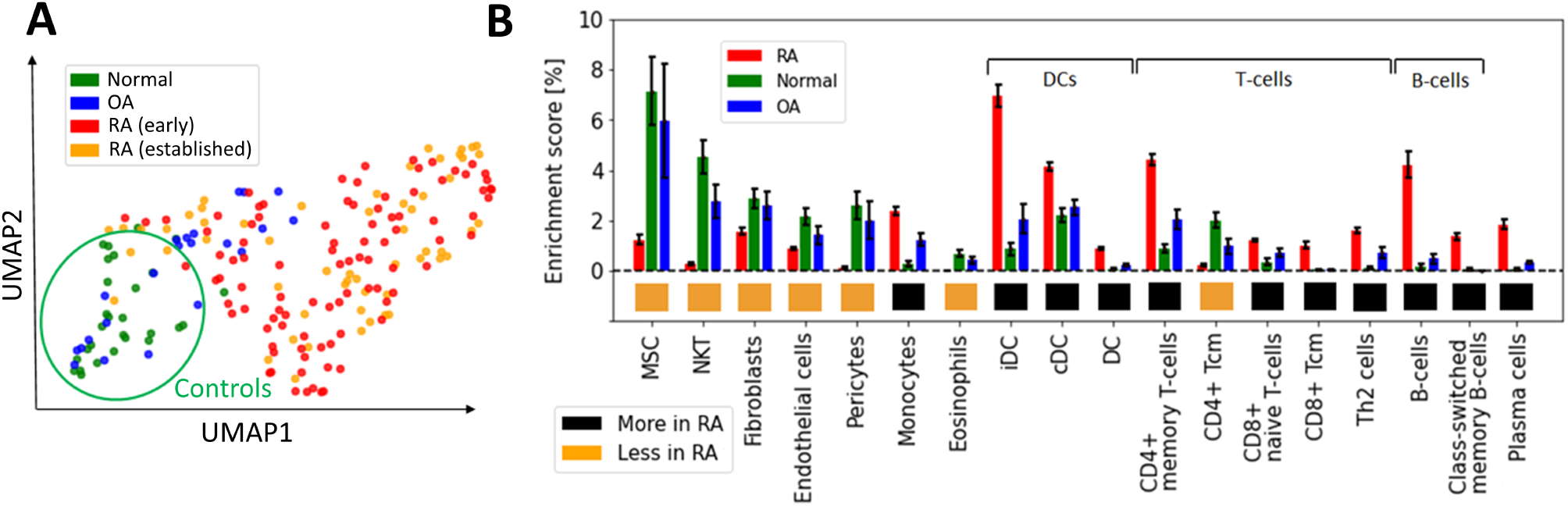
Heterogeneous cellular composition in synovial tissues. (A) UMAP representation of the gene expression data, with samples colored as a function of diagnosis. Healthy and RA biopsies are projected quite far apart from each other. (B) Cell types with significantly different enrichment scores in both RA *vs* normal and RA *vs* OA tissues (*p <* 0.05, Student’s *t*-test), ordered by average enrichment score across tissues, except for DCs, T cells, and B cells that were grouped together for visual clarity. Error bars show the 95% confidence intervals defined as 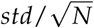. MSC: mesenchymal stem cells, DC: dendritic cells, iDC: immature dendritic cells [37], cDC: convential dentritic cells [38].

Our strategy to deconvolute synovial cellularity was able to identify populations known to be expanded in RA, such as T cells, B cells, plasma cells, dendritic cells (DC), and monocytes [15, 16, 39]. To further validate these results, we used CIBERSORT [40], an alternative deconvolution method for bulk RNA expression. Compared to xCell, CIBERSORT focuses exclusively on immune cells and employs a different algorithmic approach, namely, a linear regression based on a predefined gene expression matrix of known cell types (Supplementary Section A.2 and Supplementary Figure S2). CIBERSORT and xCell findings were similar regarding the enrichment of plasma cells, memory B cells, CD4+ memory T cells, and dentritic cells (DC), thus confirming the different synovial cellularity between RA and controls (Supplementary Table **??**).

In addition to these well-documented cell populations, xCell also identified other cell-specific traits associated with RA, such as an increased representation of both immature dendritic cells (iDC) [37] and conventional dendritic cells (cDC) [38, 41], as well as a previously unrecognized reduction of Natural killer T cells (NKT), pericytes and eosinophils, compared with controls. These analyses revealed the high variability in cellular composition within synovial tissues, which may explain a significant fraction of the gene expression variability observed among RA patients [14]. To quantify this effect, we calculated the expression variance explained by cellular composition using the R-squared score. This statistical measure represents the proportion of total gene expression variance explained by the enrichment scores estimated with xCell. xCell scores were used as inputs in a linear regression model to predict gene expression values. The R-squared score was then derived by comparing the model’s predictions with the original observations. Significantly, we found that the 18 cell types shown in Figure 1B account for 61% of the variance in gene expression (Supplementary Section A.3, Supplementary Figure S3).

### 1.2 Gene regulation in RA *vs* control samples differs widely across synovial cell types

The measured gene expression in synovial tissues is a mixture of gene expression profiles from different cell types, which complicates the task of extracting cell type-specific gene regulatory information. To address this challenge, we adjusted the gene expression values to correct for cellular composition biases using a linear model (Methods Section 2.3). The corrected gene expression data served as a basis for constructing a gene regulatory network, unbiased by cell type heterogeneity. In addition, we also leveraged cell type-specific bulk RNA-Seq data from RA and OA synovial fibroblasts, monocytes, B cells, and T cells [15] to complement our analyses of the synovial tissue biopsies, resulting in 5 independent gene expression datasets from synovial tissues (Table 1).

**Table 1:** List of datasets of RNA-Seq data from synovial tissues and bipartite networks computed from them. An individual network is computed for each RA and OA sample.

| Dataset/networks name | Reference | Method | Number of patients/networks |
| --- | --- | --- | --- |
| PANDA_Synovial_Tissue | [26] | PANDA, LIONESS | 152 RA, 22 OA |
| PANDA_FLS | [15] | PANDA, LIONESS | 18 RA, 12 OA |
| PANDA_Synovial_Monocyte | [15] | PANDA, LIONESS | 17 RA, 13 OA |
| PANDA_Synovial_B cell | [15] | PANDA, LIONESS | 8 RA, 7 OA |
| PANDA_Synovial_T cell | [15] | PANDA, LIONESS | 17 RA, 13 OA |

Focusing on the cell type-specific datasets, we used a Student’s t-test to identify the genes that were differentially expressed between RA and OA samples. Here, we considered OA samples as the control group due to the unavailability of biopsies from healthy patients. From this analysis, we obtained a differential expression score for each gene (denoted *t*_expr_). Next, we examined the correlation of these scores *t*_expr_ across each pair of cell type-specific datasets (Methods Section 2.5). The correlation among the genes’ differential expression scores was notably low (< 0.1) across all considered datasets, suggesting disparate regulatory mechanisms for each RA-associated cell type (Supplementary Figure S7).

Nonetheless, this approach failed to provide insights into the regulatory connections among the DEGs. It also did not reveal whether these connections involved TF-mediated inhibition, activation, or co-regulation. To investigate this question, we assembled gene regulatory networks of the synovial tissue and its constituent cell types by combining RA gene expression data with information about TF-binding motifs (CIS-BP [42]) and protein-protein interactions (StringDB [43]). We leveraged PANDA [44], a computational strategy designed to optimize the alignment between gene expression data and pre-existing knowledge (Methods Section 2.8), and used LIONESS [45] to estimate individual gene regulatory networks for each sample in our cohort. In these sample-specific networks, each edge connecting a TF and a target gene (TG) has an associated weight that represents the likelihood of a regulatory interaction between the TF and the TG. We leveraged this collection of networks to evaluate whether the TF-TG interaction edge weights differ significantly between the RA and control samples and to identify potential TFs that might regulate the regulatory differences (Figure 2A). More precisely, we compared the differences in edge weights between RA and OA samples using a Student’s t-test, and obtained a score *t*_edge_ for each edge TF → TG. For each cell type, we assembled all *t*_edge_ scores in a *differential GRN* (dGRN) network, which highlights the edges that are differentially regulated between RA and OA (Fig. 2A). Then, to quantify TFs regulatory function, we define a TF regulatory score (*t*_reg_) defined as the mean of the absolute values of the edge scores (|*t*_edge_~) between the TF and all its TGs (Method 2.9). As a positive (respectively negative) *t*_edge_ indicates a strong upregulation (respectively downregulation) TF-TG interaction in RA with respect to OA (control) tissues, TFs with a high *t*_reg_ are potential key regulators to explain the differential gene expression between RA and OA.

**Figure 2:**
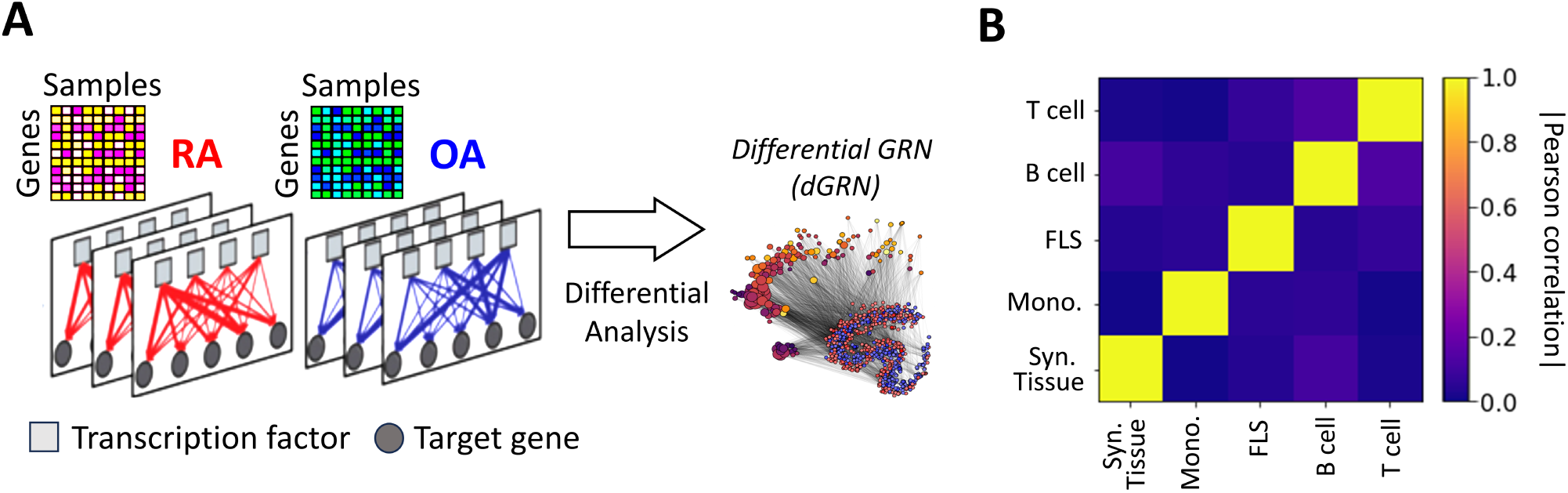
(A) Networks were inferred from the gene expression profiles of RA and OA biopsies in different cell types. Network edges are endowed with weights representing the probability of regulatory interactions between a transcription factor (TF) and a target gene (TG). The analysis of differences in edge weights between RA and OA facilitated the construction of a differential GRN (dGRN) for each cell type. The dGRN was used to compute a regulatory score for each TF in each cell type. (B) Heatmap of the Pearson correlation between the TF regulatory scores in each tissue type, i.e. synovial tissue, monocyte, FLS, B cell, and T cell).

Table 2 lists the top 10 TFs ranked by their regulatory scores for each synovial tissue cell type. Interestingly, several TFs, including RFX5, CEBPZ, SCRT1, and MXI1, are in the top 10 in more than one cell type, suggesting that these TF are broader key regulators in synovial tissues. We examined the correlation between the TF scores (*t*_reg_) on each pair of networks and obtained similar results to those previously observed with the cell-specific gene expression signatures, i.e. a low overall correlation (*<* 0.1) indicating distinct regulatory mechanisms across cell types (Figure 2B).

**Table 2:** Top 10 ranked TF regulators in each synovial cell type, along with their Z-scores in parentheses. TFs found in the top 10 in more than one network are highlighted in bold. Z-statistics for all TFs in each network are available in Supplementary Data 1).

| Rank | Syn. Tissue | Syn. Monocyte | Syn. Fibroblast (FLS) | Syn. B cell | Syn. T cell |
| --- | --- | --- | --- | --- | --- |
| 1 | ZNF282 (7.01) | NFE2 (5.71) | RORC (3.39) | JUN (6.59) | HAND1 (5.85) |
| 2 | SCRT2 (6.30) | **CEBPZ** (5.63) | NKX2-1 (3.37) | STAT5B (4.89) | REST (4.97) |
| 3 | NR6A1 (4.06) | IRF8 (5.59) | HOXA1 (3.20) | CTCF (4.04) | ELF4 (4.59) |
| 4 | **SCRT1** (3.70) | PBX3 (5.60) | ETV2 (2.67) | TCF3 (3.45) | EHF (4.21) |
| 5 | FOSL2 (3.11) | NFYA (5.49) | MITF (2.64) | JUND (3.37) | NR4A1 (3.36) |
| 6 | **RFX5** (2.52) | **RFX5** (4.62) | SIX5 (2.61) | NKX2-3 (3.20) | **SCRT1** (3.32) |
| 7 | **MXI1** (2.41) | IRF9 (3.97) | ELF5 (2.46) | KLF13 (3.03) | NRF1 (3.19) |
| 8 | ZBTB33 (2.29) | **MXI1** (3.32) | ZBTB1 (2.42) | PBX3 (2.69) | EPAS1 (2.94) |
| 9 | JUNB (2.07) | IRF4 (3.21) | FOXC2 (2.27) | TWIST1 (2.69) | NFYB (2.81) |
| 10 | RELA (2.05) | ARID2 (2.88) | TBX4 (2.23) | FOSB (2.52) | **CEBPZ** (2.77) |

For additional insight into the pathways involved in RA in each cell type, we ran a pathway enrichment analysis on the major TF regulators using a collection of pathways compiled from the Gene Ontology (GO) [46], KEGG [47], and Reactome [48] databases. First, we selected the TFs with a *t*_reg_ score higher than one standard deviation above the mean (Z-statistics > 1). This led to the selection of between 40 and 90 TFs per tissue and cell type (Figure S8A). Interestingly, the overlap between the selected TFs was low, and only 7 TFs were shared by 3 cell types or more (Figure S8B). Because the enrichments were run with TFs exclusively, we removed any terms associated with RNA and DNA transcription, as these are ubiquitous processes and not likely to be RA-specific. After this filtering, we obtained between 60 and 120 significantly enriched pathways (i.e. *p*_adj_ < 0.05) for each cell type (Figure S9A). In Figure 3, we show the 10 most significant pathways for each cell type. As expected, well-established RA pathways are evident across multiple cell types, such as TNF [49], IRF [50] and ATF6 [51], along with pathways associated with osteoclast differentiation [52] and T helper (Th) cell differentiation [53]. Our analysis also identified less commonly documented pathways in the context of RA, such as those associated with RUNX1, found in monocytes and fibroblasts, and HOX, predominantly in fibroblasts and T cells.

**Figure 3:**
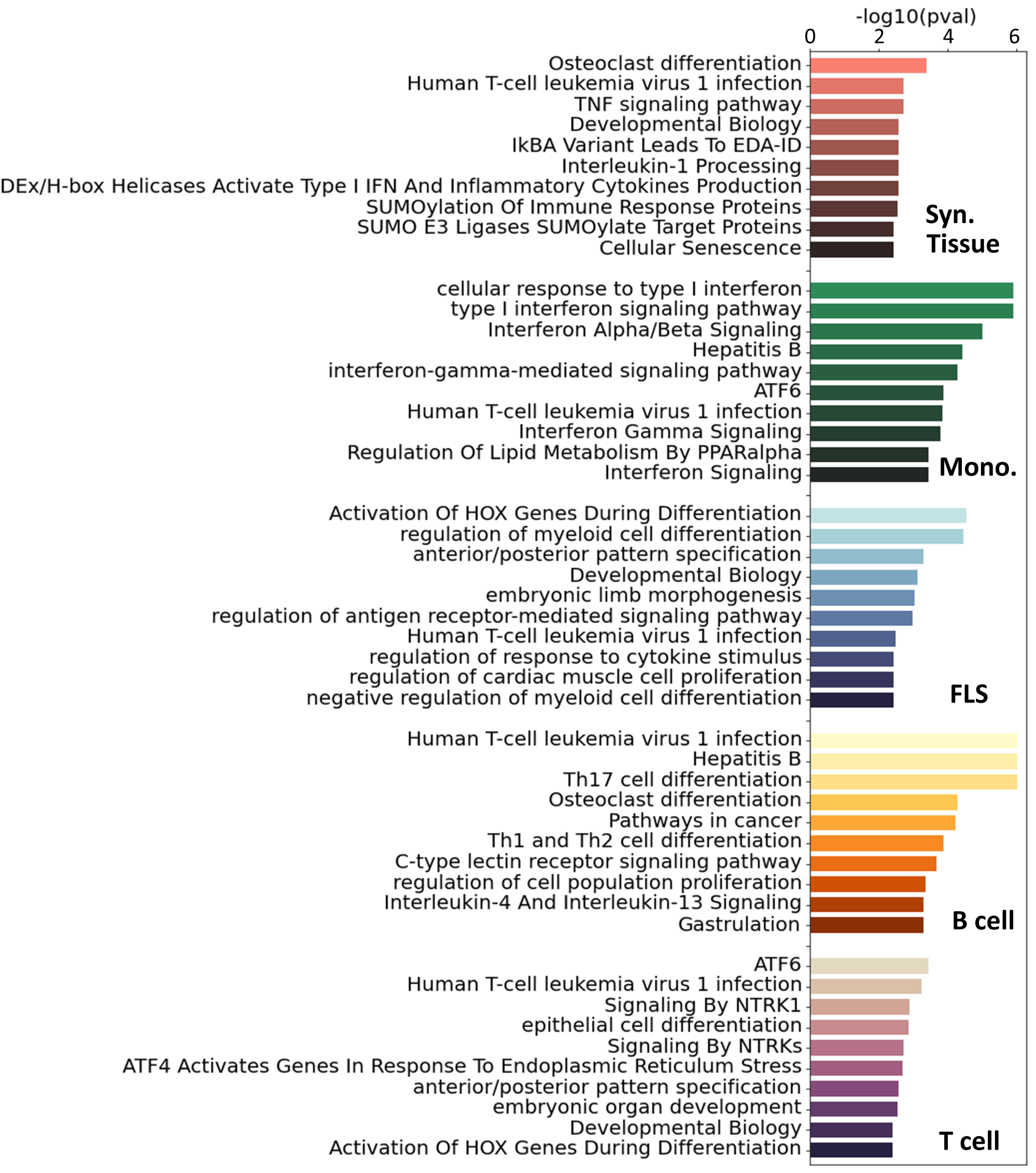
Top 10 significant pathways enriched on the main TFs involved in RA for each specific cell type. Pathways were compiled from GO [46], KEGG [47] and Reactome [48]. These pathways were ranked based on adjusted *p*-values.

To get a more complete overview of the ~400 significant pathways that were identified across the three pathway databases (GO [46], KEGG [47], Reactome [48]), we split each term by words and counted the accumulated word count found in each cell type (Figure S9B). We identified, among others, T cells, alpha-beta, myeloids, and B cells, as words that appear consistently across cell types, specically T cells had 24 occurrences, “*α*/*β* signaling” 9 occurrences, myeloid cells 8 occurrences and B cells 7 occurrences.

### 1.3 A subset of RA key driver genes are consistently identified across cell types and tissues

The analysis of the sample-specific networks identified a list of candidate master regulator TFs in RA, and generated detailed statistics about the regulatory function of these TFs and their TGs within each cell type Supplementary Data 1) However, different network inference methods exhibit considerable variability in their inferred networks [54], typically due to the varying algorithmic assumptions and limited sample sizes. In our study, sample sizes were small in all considered cases, ranging from 15 to 40 samples for the cell type-specific networks. Hence, relying on the predictions of a single computational method might lack the robustness required to identify promising therapeutic targets. To increase our confidence in the identified RA regulators, we augmented our study by incorporating a selection of pre-existing literature-derived networks, which also included edge weights as a metric for assessing the confidence of the interactions between nodes. These include (i) RIMBANET [55], a probabilistic causal network reconstruction approach that integrates multiple data types, including metabolite concentration, RNA expression, DNA variation, DNA–protein binding, protein–metabolite interaction, and protein–protein interaction data; (ii) StringDB [43], a database of known and predicted protein–protein interactions from numerous sources, including experimental data, computational prediction methods, and public text collections; (iii) GIANT [56], a collection of networks that accurately capture tissue-specific and cell type-specific functional interactions. As RA is an autoimmune disease, we selected networks computed from immune-related tissues (including lymph nodes, spleen, tonsils, and blood). Additionally, when available, we extracted networks associated with cell types present in different proportions in RA vs control patients (Section 1.1). 14 additional networks were collected for our analysis, as detailed in the Supplementary Table S2.

While these networks recapitulate general immune knowledge derived from various data types, they are not specific to synovial tissues. Therefore, they are unable to discern RA-specific relationships between TFs and TGs as effectively as the PANDA framework does. We hence designed a different approach based on the *key driver analysis* (KDA) [57], a computational pipeline to uncover major disease-associated regulators or causative hubs in a biological network (Methods Section 2.7). Briefly, genes exhibiting more connections to RA-associated genes than expected by random chance were considered potential drivers (Figure 4Af). Using a list of RA-associated signatures as a starting point, we identified potential *key drivers genes* (KDGs) linked to these signatures via network edges. To mitigate potential network size bias in the identification of KDGs, we only considered the top 1 million edges in each network. Note that a fully connected network of ~40k genes contains more than 1 billion edges, and hence, the selected edges roughly represent the top 0.1% network interactions.

**Figure 4:**
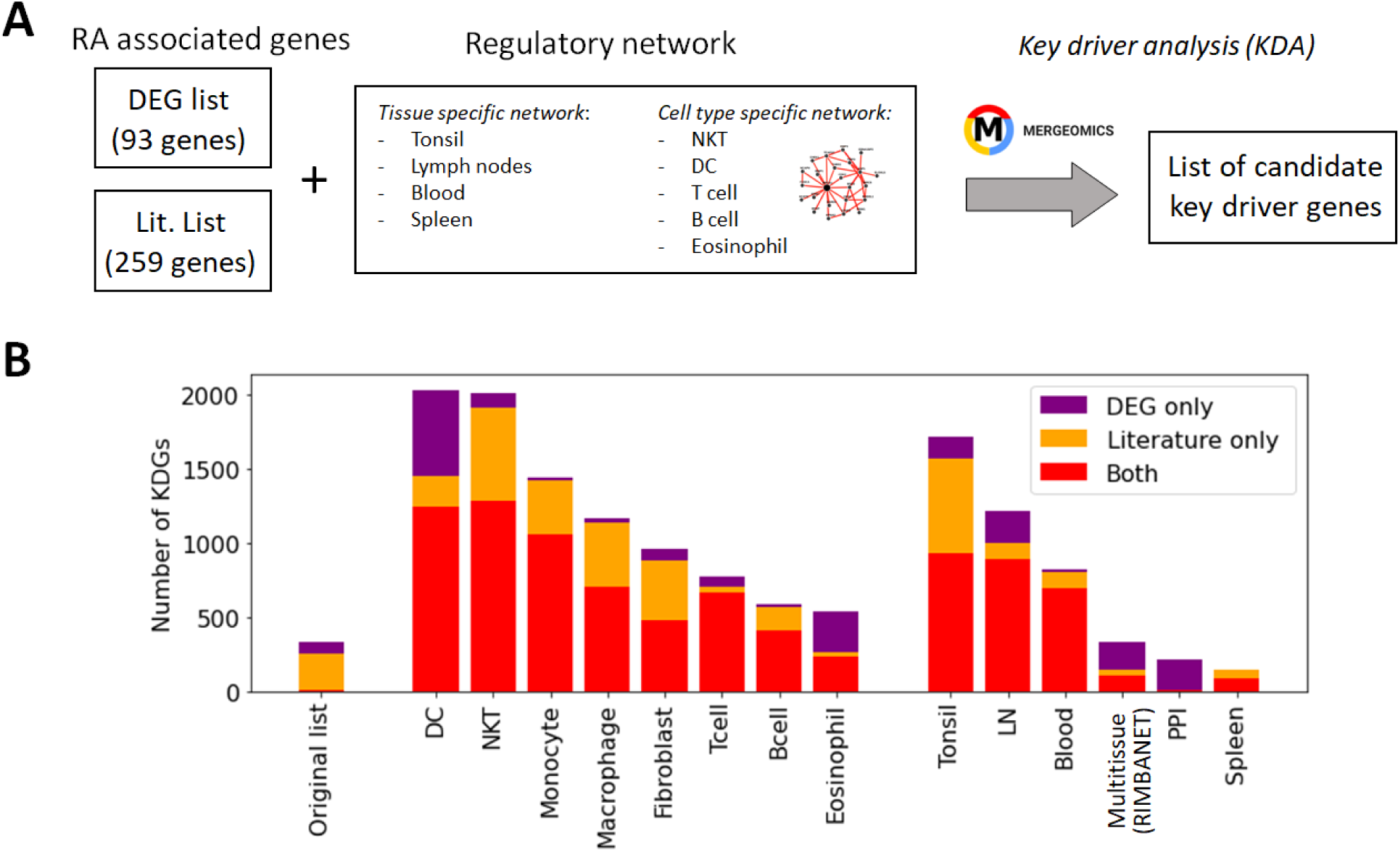
Identification of key driver genes (KDGs). (A) Within the Mergeomics framework [68], RA signature genes are leveraged to test the significance of each gene node within a given network. We performed the analysis with 2 different RA signature gene sets (*DEG list* and *Literature list*) and 10 networks. (B) Number of obtained KDGs for each tested network using the *DEG* and *Literature lists*. While the overlap between the RA signature gene sets is small, we observe a high overlap (in red) between the KDGs inferred from both lists across different cell types. KDGs for all networks are provided in Supplementary Data 2). DC:dendritic cells; LN: lymph node; PPI: Protein-Protein interaction network.

KDA analysis requires the definition of RA-associated signatures, i.e. lists of genes associated with the disease. A common practice is to create gene signatures from DEGs [23, 25]. However, these signatures are likely to be biased by heterogeneity in cellular composition and might include bystander genes that are not directly linked to the pathogenesis of the disease. Hence, we constructed two independent lists for KDA analysis. The first list exploited prior meta-studies and datasets [58, 59, 60, 12, 13] from which a list of DEGs was extracted (Supplementary Section B.1). The second signature combined known RA-associated genes from the literature, including GWAS [61, 10, 62], knowledge-based datasets [63, 64, 65], and known drug targets [66, 67] (Methods Section 2.6). To summarize, we performed KDA on 14 different networks (Table S1), with two RA-associated gene signatures, which we refer to as the *DEG list* (93 genes) and the *gene literature list* (259 genes). Interestingly, the overlap between these databases was moderate (~2000 genes in total after combining all databases). Additional information is provided in Table 3 and Supplementary Section B.2.

**Table 3:** Genes associated with RA and those identified with key driver gene (KDG) analyses. (A) Number of genes utilized for KDA analysis in two different lists. The intersection is calculated as *A B* /*min*(|*A*, *B|*). (B) Mean number of genes designated as KDG following KDA in various networks from the two lists.

| A. RA associated genes |  |  | B. Obtained KDGs (averg. over networks) |  |  |
| --- | --- | --- | --- | --- | --- |
| Lit. list | DEG list | Overlap | Lit. KDG | DEG KDG | Overlap |
| 259 | 93 | 10% | 741 | 827 | 90% |

While the overlap between the *literature list* and the *DEG list* was quite small, i.e. overlap = |*A*⋂ *B* /*min*(|*A|*,|*B|*) = 9/93 genes 10%, they converge to a similar set of driver genes after KDA in all analyzed networks (average overlap of 90%, Table 3 & red bars in Figure 4B). This suggests that there are common drivers behind the set of DEGs and the set of known RA-associated genes reported in the literature. The number of identified KDGs, even when derived from the same lists of RA-associated genes, varied significantly across the different cell type-specific networks used in the analysis. The highest number of KDGs was found in DC and NKT, and tonsils & lymph nodes, among the various tissue types (all identified KDGs in each network are available in Supplementary Data 2). This suggests that crucial RA regulation occurs in these cell types as well as within these tissues. We also found that several genes were consistently identified as key drivers across most networks. More precisely, more than 500 genes were found in more than half of the tested networks (Supplementary Section C and Supplementary Figure S6), and the top 20 genes were found in 75% of the networks (Table 4). Among them, there are several that were already included in our *DEG* or *Literature* lists (HLA and IL2 variants, CCL5, PSMB8, CTSH), but also some that were not (PTPN6, SRGN, GBP1, LCP2, GLIPR1, CTSS, CTSH, CASP1, CD44). Importantly, the majority of genes in our top 20 list had been previously characterized in the context of RA (Table 4), thus providing further validation for our discovery approach.

**Table 4:** The top 20 KDGs identified using the KDA with both the *DEG* and *Literature lists* are presented below. Genes also identified in GWAS are highlighted in bold. The complete list of the top 100 KDGs is available in Supplementary Section C. The left (right) number corresponds to the number of networks the TF was identified as a KDG with the literature (DEG) list (adjusted *p*_val_ *<* 0.05).

| Gene | KDA<sup>†</sup><br>(Lit-DEG) | Reference | Gene | KDA<sup>†</sup><br>(Lit-DEG) | Reference |
| --- | --- | --- | --- | --- | --- |
| PTPN6 | 12 - 12 | [69] | CASP1 | 11 - 11 | [70] |
| HLA-E | 12 - 11 | [71] | **HLA-B** | 12 - 10 | [71] |
| HLA-F | 12 - 11 | [71] | **HLA-C** | 12 - 10 | [71] |
| GBP1 | 12 - 11 | [72] | CD44 | 12 - 10 | [73] |
| LCP2 | 12 - 11 | [74] | IFI30 | 12 - 10 | None |
| GLIPR1 | 11 - 11 | None | TNFAIP8 | 12 - 10 | [75] |
| **HLA-A** | 11 - 11 | [71] | CCL5 | 12 - 10 | [76] |
| CTSS | 11 - 11 | [77] | **PSMB8** | 12 - 10 | [78] |
| SRGN | 11 - 11 | None | TAP1 | 12 - 10 | [79] |
| CTSH | 11 - 11 | None | ICAM1 | 12 - 10 | [80] |

### 1.4 Combining the KDA and PANDA analysis

For each cell type, we compiled a list of key regulators by retaining TFs that met the following criteria: (i) Their regulatory scores (*t*_reg_), computed with the PANDA network specific to each cell type, were at least one standard deviation above the mean score of all TFs (Z-statistics > 1); and (ii) they were identified as KDGs by both the *DEG* and *Literature lists* in at least one of the 14 networks. We obtained between 10 and 18 TFs for the different cell types. The full list is available in Table 5. Interestingly, while the majority of key regulators were specific to individual cell types, we found that several TFs, such as RFX5, RELA, FOS, HIVEP1, IRF9, MITF, ETV7, FOSL1, FOSB, KLF2, and ELF4, were identified as key regulators in two or more cell types.

**Table 5:** Key transcription factors (TFs) implicated in the regulation of RA identified in our analyses (Z-statistics > 1), ordered from highest to lowest Z-statistics for the different cell types. TFs identified in more than one cell type are highlighted in bold.

| Cell type | ( # of TFs) | List of TF drivers (Ranked from highest to lowest Z-statistics) |
| --- | --- | --- |
| Syn. Tissue | (10) | FOSL2, **RFX5**, **RELA**, JUNB, **FOS**, MYBL1, NFKB2, ZNF274, **HIVEP1**, HIVEP2 |
| Syn. Monocyte | (11) | IRF8, **RFX5**, **IRF9**, IRF2, CREB5, IRF3, XBP1, STAT2, EOMES, BCL11A, SPI1 |
| Syn. Fibroblast | (18) | **MITF**, CBFB, IKZF1, HLX, BACH1, **FOS**, **ETV7**, **RFX5**, HIF1A, TGIF1, **ELF4**, RUNX1, NFATC1, IRF7, CREM, **FOSL1**, FLI1, ELF1 |
| Syn. B cell | (19) | STAT5B, JUND, KLF13, **FOSB**, **FOSL1**, **KLF2**, PLAGL1, EGR3, CEBPD, TCF7, RELB, HBP1, RUNX3, BATF3, RORA, STAT6, EGR2, BHLHE40, CIC |
| Syn. T cell | (15) | **ELF4**, NR4A1, ETV6, **MITF**, ETS2, **ETV7**, MSC, **FOSB**, **RELA**, **RFX5**, **KLF2**, REL, ATF4, **HIVEP1**, VENTX, **IRF9** |

To evaluate the agreement between the two methods we used, namely KDA and PANDA, we examined whether TFs identified as key drivers in one of the 14 networks exhibited, on average, higher PANDA scores (*t*_reg_) than other TFs. Interestingly, we found that, on average, the TFs identified by KDA in any of the tested networks had a significantly higher regulatory score in the PANDA networks than the genes not identified by KDA (*p* = 1 × 10^−5^, Wilcoxon Signed-Rank Test). Figure 5 illustrates the positive relationship between the KDA and PANDA scores, where KDG TFs identified with KDA typically exhibit higher PANDA scores in at least one cell type.

**Figure 5:**
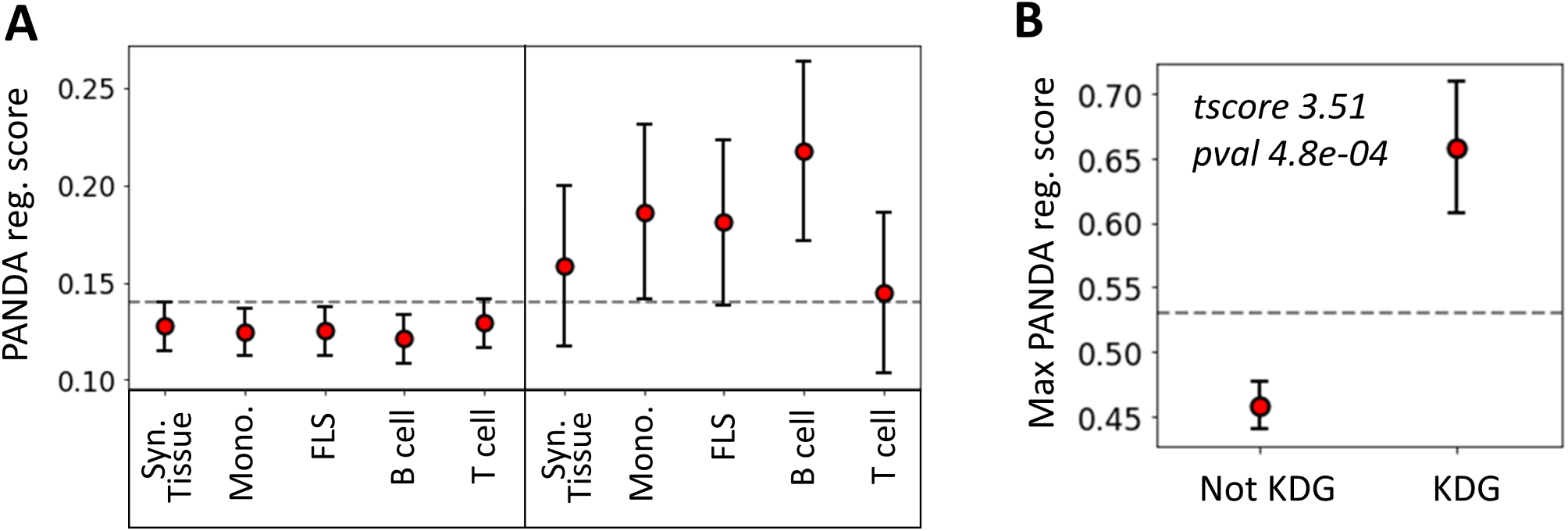
Relationship between key driver analysis (KDA) and PANDA network-based TF analysis. (A) Average PANDA score of TFs identified as key driver genes (KDA score > 2) and not identified by the KDA analysis (KDA score ≤ 2) for each tissue type. (B) Max PANDA score (defined as the maximum PANDA score across all cell types) for both key driver TFs (KDA score > 2) and non-key drivers (KDA score ≤ 2). The gray line represents the expected score if <u>al</u>l TFs were randomly scored, and the error bars correspond to the 95% confidence intervals, defined as 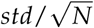.

### 1.5 Comparing the TF-TF co-regulation network across cell types

While we have identified a list of key TF regulators, it remained uncertain whether these regulators collectively controlled the same genes and phenotypes, or whether they independently regulate distinct targets. Clarifying the potential co-regulatory role of these TFs might open the door to combined therapeutic strategies targeting multiple TFs simultaneously. To investigate key RA driver co-regulation within each cell type, we computed the Pearson correlation between the differential edge weight *t*_edge_ of the common TGs between two TFs.

To maintain consistency across all cell types, we selected the top 30 key driver TFs ranked by their Z-statistics per tissues type, and computed pairwise correlations between all key driver TF-TF pairs in these subsets. Then, in each cell type, we employed a hierarchical clustering algorithm with a correlation threshold of 0.5 to cluster these 30 TFs based on their pairwise co-regulation patterns (Figure 6A).

**Figure 6:**
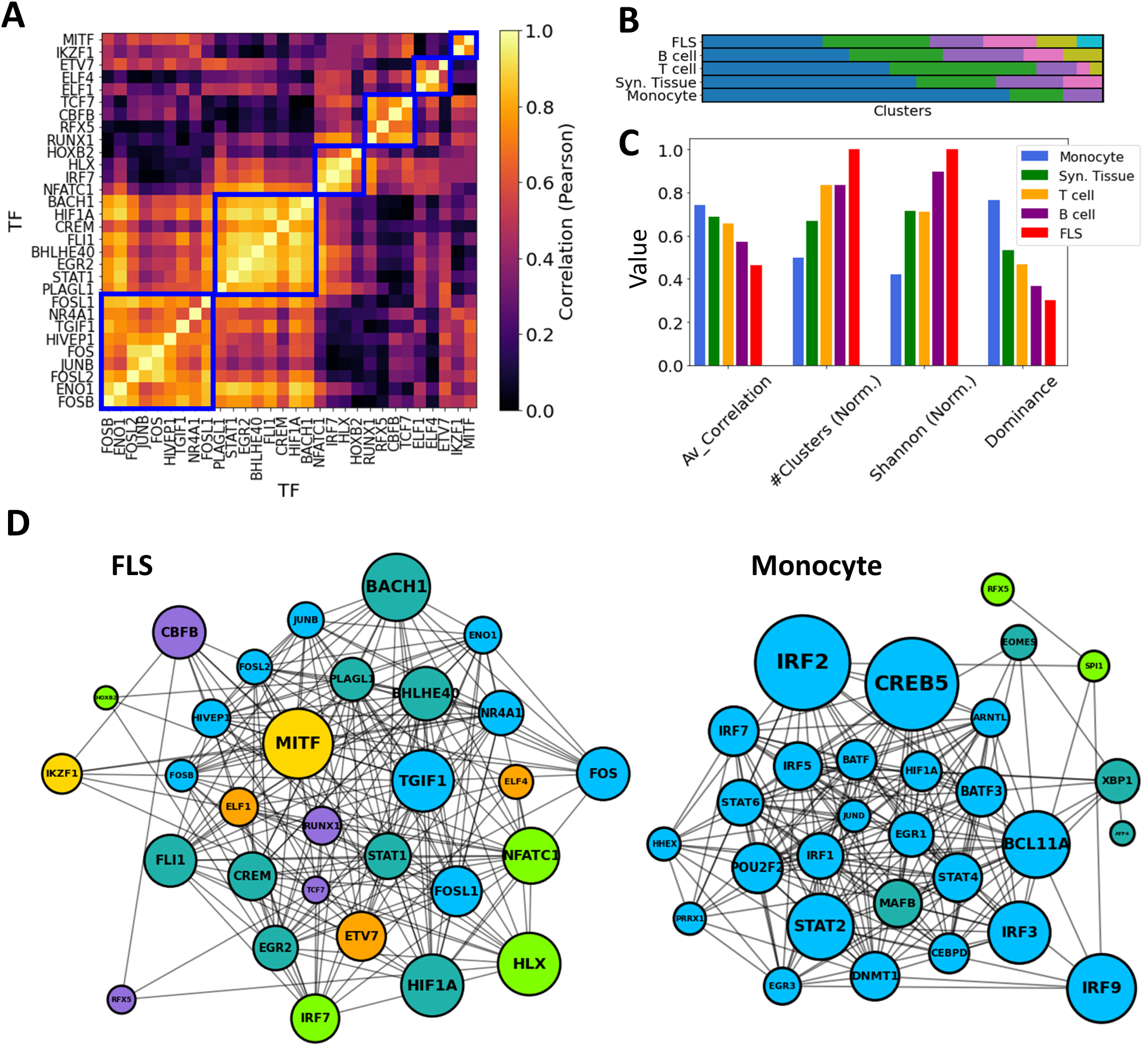
TF-TF RA co-regulation network in each cell type. (A) Pairwise TF-TF co-regulation heatmap in FLS, quantified in terms of the Pearson correlation between the differential edge weight *t*_edge_ to their common target genes (Method 2.10). A hierarchical clustering approach is used to group them into clusters (depicted with a blue square). (B) The resulting clusters are visualized as a bar plot, with populations from each cluster depicted in distinct colors. (C) The networks are characterized by the average correlation of their edges, their number of clusters, and other diversity metrics (Shanon Entropy, dominance). These metrics are plotted next to each other for each cell type. For visual clarity, some metrics were normalized by their maximum values. (D) Networks show the main TFs involved in FLS and monocyte RA regulation. Edges indicate correlations exceeding the median co-regulatory scores (Methods Section 2.10). Node sizes are proportional to the node degree times TF’s regulatory score, (*t*_reg_). Node colors indicate the different co-regulatory clusters. Networks and pairwise correlation matrices are provided for all cell types in Supplementary Figure S10.

Significantly, monocytes exhibited a notably strong overall correlation of their TF drivers, with an average pairwise correlation coefficient of 0.77. This suggests a coordinated regulation of the observed transcriptomic distinctions between RA and OA FLS. Conversely, FLS displayed an average pairwise correlation coefficient of 0.46, and their regulators were divided into 7 clusters with minimal correlation among them. This suggests the co-existence of multiple independent FLS regulation clusters targeting different genes contributing to RA pathogenesis. T cell and B cell co-regulation clusters had an average correlation coefficient of 0.69 and 0.56, respectively. These findings are illustrated in Figure 6B, where distinct cluster populations across cell types can be observed. To quantify these disparities, we computed various diversity indices, such as dominance and Shannon entropy [81, 82] (Methods Section 2.11), and observed substantial differences across cell types in all metrics. This divergence is also evident in Figure 6C & D, where the generated TF-TF networks exhibit distinct co-regulated clusters in the case of FLS, in contrast to monocytes, which are predominantly regulated by a single co-regulatory cluster.

## Discussion

RA is a common autoimmune and inflammatory disease that affects nearly 1% of the population [3]. Despite significant advances in treatments targeting different aspects of the immune response over the last two decades, achieving sustained disease remission remains uncommon [83]. A small percentage of patients may achieve complete disease control on one type of therapy, yet predicting treatment responses remains challenging. This variability in treatment efficacy could be partly due to the diverse genetic factors influencing the disease, with certain genes playing more prominent roles in some patients. Additionally, variations in the disease itself and differences in individual patients’ synovial tissue cell composition and activation states may contribute to this variability. A third and important missing component is the role of central genes in regulating transcriptional networks in different cells, along with their interaction and co-regulation. In this study, we combined recently published RNA-Seq data with innovative bioinformatics and analytical techniques. We identified previously uncharacterized transcriptional networks, along with their essential driver genes and TFs in synovial tissues and synovial cells from RA patients.

Extensive new RNA-seq databases and studies have significantly deepened our understanding of disease processes in RA. Nevertheless, conventional bioinformatics workflows for data-driven discovery, including GWAS, DEG analyses, and cohort-averaged GRNs, are not without challenges. For example, many GWAS loci are situated outside protein-coding regions, which complicates their functional interpretation [84]. Additionally, DEGs can identify many genes that are not causally associated with the disease [85], and GRNs inferred from heterogeneous samples fail to discriminate cell type-specific regulatory mechanisms. Gene regulatory mechanisms vary significantly across cell types in both health and disease. Elucidating cell type-specific pathogenic mechanisms associated with major diseases can significantly facilitate the development of novel therapeutics, with the potential to target specific networks, pathways and cell types, with reduced risk for side effects. However, the identification of cell type-specific regulatory processes remains a challenge [84]. In this context, we have developed a novel computational pipeline to identify key drivers underlying RA pathogenesis in synovial tissues. A key aspect of our analysis pipeline is the inference of sample-specific GRNs, as opposed to the more commonly used cohort-specific GRNs. This approach offers significant advantages: it reveals the regulatory mechanisms associated with individual samples and facilitates the use of statistical techniques to compare network properties across samples and phenotypic groups [86]. Our approach also enabled us to rank TFs based on their contribution to phenotypic disease differences.

In the context of RA, our analysis revealed that conventional DEG analyses of synovial tissues were heavily confounded by the heterogeneous cellular composition across tissues. Indeed, we discovered that 60% of the variability in gene expression could be attributed to varying cell type proportions rather than actual differences in tissue gene regulation. Interestingly, biopsies from early and established RA had similar gene expression signatures and cellularity, suggesting that similar cell types and processes regulate disease all through the different stages of progression and, therefore, therapies can be effective throughout the disease course. Among the overrepresented cell types in RA tissues compared with control samples were DC, CD4+ memory T cells and B cells. Conversely, NKT cells emerged as the most statistically significant underrepresented cell type.

The observed reduced numbers of NKT cells in synovial tissues of RA patients suggested either impaired differentiation or impaired tissue migration or chemotaxis. Numbers of NKT cells were previously reported to be decreased in the peripheral blood of RA patients [87]. NKT cells typically accumulate in the liver and move into tissues in response to chemotactic factors such as CCL5, CXCL16 and others, which are known to be expressed in the RA synovial tissues [88]. NKT cells express an invariant TCR that recognizes glycolipids presented by CD1d [89]. Synthetic versions of these glycolipids have been used to successfully treat arthritis in rodent models [90, 91]. Additionally, recently discovered and naturally-occurring glycolipids produced by Bacteroides fragilis were shown to induce the differentiation and activation of NKT cells [92, 93]. Interestingly, Bacteroides fragilis is a bacterial species commonly depleted in the intestinal microbiome of RA patients, raising the possibility of a connection between the intestinal microbiome and the reduced numbers of NKT cells in synovial tissues and blood of RA patients [94]. Functionally, NKT cells can produce IL-10 to suppress immune responses, and also can inhibit autoreactive B cells [95], which are expanded in RA synovial tissues and have central in disease. Therefore, the reduced NKT numbers could favor an expansion of the autoimmune and inflammatory response in the synovial tissues[96].

Another interesting cell population that emerged from our analysis were eosinophils, which were almost absent in RA tissues compared to OA and healthy controls. Interestingly, eosinophils with a regulatory phenotype were recently reported in the synovial tissues of RA patients in remission [97], suggesting that their presence may help control disease. Likewise, eosinophil activation can suppress inflammation in arthritis in rodent models [98].

To elucidate the potential role of the RA-associated candidate genes identified in our analyses, we utilized previously published cell type-specific gene regulatory networks [56]. Namely, we identified genes associated with published RA signatures and used them in the key driver analysis (KDA) [57] framework. To increase the robustness of the method, we ran the analysis using two independent sets of RA-associated signatures with low overlap. The first gene set was compiled from online databases, including GWAS, knowledge-based and drug targets databases. The second gene set was developed with a DEG meta-analysis. Interestingly, numerous major regulators were consistently identified across diverse cell types and tissues. Several of these genes have been previously documented in the literature, highlighting the robustness of our methodology.

Our analyses identified new TF implicated in the regulation of RA synovial tissue gene expression, and more precisely implicating TF in cell specific gene regulatory networks (GRNs). As expected, regulatory interactions showed significant variability across cell types. Both the gene expression profiles and the regulatory edge weights of the cell type-specific GRNs showed minimal correlation. Certain cell types, such as FLS and B cells, were governed by multiple independent co-regulatory clusters, while the TF drivers in monocytes collectively controlled the regulatory distinctions between RA and OA. For example, CEBPZ was implicated in monocyte networks, SCRT1 in global synovial tissue and T cell networks, and RFX5 in global synovial tissue and monocyte cell GRNs. Most of the TF were specific to a cell type with IRF4 and IRF9 for monocyte, RORC and HOXA1 for FLS, JUN and STAT5B for B cells and ELF4 and HAND1 for T cell networks. Global synovial tissue and cell specific key drivers such as ELF4, FOSL1, FOSL2, HIVEP1, IRF9, KLF2, MITF, and RFX5 and were identified, several for the first time in RA. We also identified a major TF-TF co-regulation in the synovial tissue and synovial cells from RA patients, highlighting the complexities involved in cell regulation. It is conceivable that such networks and their dominant role in disease pathogenesis vary from patient to patient, which might help explain highly variable patient response to different treatments. But our analyses point to the likely relevant central target driving each network and may help point to new target for treatment.

Several of the KDG and TF have not been previously implicated in RA pathogenesis and their discovery opens new possibilities for studies and drug targeting. For example, SCRT1 (scratch family transcriptional repressor 1) is a recently discovered TF that has been implicated in pancreatic islet cell proliferation [99] and cancer proliferation and metastasis [100]. To our knowledge this is the first time that SCRT1 is implicated in the regulation of an inflammatory and autoimmune disease. RFX5 (regulatory factor X5) was only recently implicated in the regulation of synovial macrophage metabolism and survival [101]. Our findings suggest that this TF has a major role not only on monocyte regulatory networks but also in the synovial tissues in general. We also detected homeobox genes (NKX2-1, NK2 homeobox 1, and HOXA1) among the top KDG in RA FLS, including an overrepresentation of HOX genes pathway genes. MITF (melanocyte inducing transcription factor) is another new FLS and T-cell KDG identified in this study. MITF was previously implicated in osteoclast differentiation and function [102] and also recently shown to mediate T-cell maturation [103]. However, this is the first time that MITF is implicated in FLS and T-cell transcriptomic networks.

In conclusion, we used a robust and innovative combination of computational strategies to identify new KDG, TF and GRNs in RA synovial tissues and synovial cells. These discoveries open new possibilities for experimental validation and discovery in cells or in vivo studies [104]. Additionally, our findings should help understand the role that cellularity and the multiple pathways involved in cell regulation and co-regulations have in disease, and potentially in patient response to treatment. Unfortunately, RNA-Seq profiles of several other important cell types that significantly contribute to the heterogeneity of synovial tissues, such as NKT cells, DCs, eosinophils, or pericytes, are unavailable, and thus their GRNs could not be constructed. As such, the acquisition of additional gene expression data on these cell types could further identify candidate genes for further analyses and consideration for therapeutic development [16].

## 2 Methods

### 2.1 Gene expression data and normalization

We used two public datasets. The first one, a bulk RNA-Seq study of synovial biopsies (GSE89408) [26, 105], comprises gene expression profiles spanning over 25k genes across 28 healthy samples, 152 RA, and 22 OA patients. The second dataset is a bulk RNA-Seq study of synovial tissues including 18 RA patients with RA 13 OA patients used as controls from the Accelerating Medicines Partnership (AMP) Phase I project [15]. The first dataset is relatively bigger than the second and other publicly accessible synovial tissue studies. Moreover, it has the advantage of including both healthy and OA control groups. Also, focusing on a single big dataset, rather than combining several smaller ones, removes the batch effect bias.

All data underwent scaling normalization [106] to remove potential biases of other experimental artifacts across samples. The underlying assumption is that any sample-specific bias, such as variations in capture or amplification efficiency, uniformly scales the expected mean count for each gene. As the size factor for each sample represents the estimated relative bias in that sample, dividing its counts by its size factor should mitigate this bias.

### 2.2 Estimation of cellular compositions in synovial tissues

The cell compositions in synovial tissues were estimated with xCell [36], a machine learning framework trained using the profiles of 64 immune and stroma cell datasets. xCell takes the gene expression count as input and generates enrichment scores. Briefly, the xCell score measures the enrichment of genes specific to each cell type and is further adjusts for correlations among closely related cell types. The resulting enrichment scores are normalized to unity to enable consistent comparisons across samples.

### 2.3 Correction for cellular composition in synovial tissues

Our analysis revealed that cellular composition variation in synovial tissues accounted for a significant portion (61%) of gene expression variability. To differentiate gene expression variability arising from actual molecular state changes in cells from those due to compositional shifts, we adjusted the gene expression profiles for these covariates. To prevent over-correction, we corrected only the 18 cell types that exhibited significant differences in both RA *vs* normal and RA *vs* OA comparisons (Student’s *t*-test with *p <* 0.05 after Benjamini-Hochberg correction [29]).

For each gene *k* in sample *s*, we performed a linear regression analysis using the proportion of cell types *c_i_* on that sample *s* as covariates, i.e. *x_ci_*_,*s*_. Mathematically, this is expressed as:

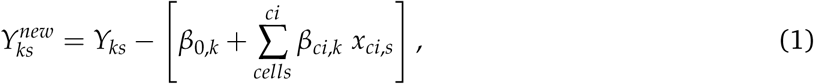

where the *β*s are regression coefficients derived using a least-square fit. We then used the residuals from this regression, (*Y_ks_*), as the actual gene expression value in our analyses [107]. Namely, we utilized these adjusted gene expressions for differential gene expression (DEG) analysis and to assemble the RA synovial network.

### 2.4 Differentially expressed genes (DEGs)

After correcting the gene expression data for cell composition, we defined differentially expressed genes (DEGs) as genes with a *p*-value below 0.05 in a *t*-test comparing the RA and control groups, after applying the Benjamini & Hochberg method [29] to control the False Discovery Rate (FDR) at 0.05. This approach ensures that, on average, only 5% of the identified DEGs are false positives, offering a robust balance against multiple testing errors.

Because the DEG analysis relies on an error-prone cell composition correction of the synovial tissues, we combined our DEGs with several meta-analyses [59, 60, 12, 13] from synovial tissues to increase the DEG analysis robustness. Genes that were identified in at least two of the lists above (either our DEGs or from one of the meta-studies) were kept as the *gene DEG list* (93 genes).

### 2.5 Correlation of gene expression across cell types

For a given gene *g* ∈ *G*, we performed a Student *t*-test to compare the difference in expression values between RA and the control group within a specific cell type *C*. From this test, we obtained its t-statistic, denoted as *t*_expr_(*g*, *C*). Next, we calculate the correlation of this score across cell type pairs (*C*1, *C*2), as follows:

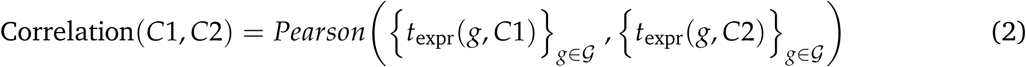

### 2.6 Genes associated with the susceptibility to RA

We performed an extensive literature review to aggregate known genes associated with RA from different contexts. Recent GWAS data, highlighting genetic risk factors for RA, were collected from publicly available studies [61, 10] and the GWASdb SNP-Disease Associations database [62]. Various genes associated with RA were obtained from publicly-available databases DISEASES [64], DisGeNet [63] and the Comparative Toxicogenomics Database (CTD) [65]. Briefly, the CTD score, ranging from 0 to 100, measures the deviation of a gene’s connectivity in the CTD chemical-RA network from that of a random network. (see http://ctdbase.org/help/diseaseGeneDetailHelp.jsp for additional details). Among the ~25k genes in the database, only 175 (less than 1%) had a CTD score higher than 50. We collected drug targets either already on the market or undergoing clinical trial [66] as well as from the DrugBank database [67] (https://go.drugbank.com/categories/DBCAT003604). Genes that were identified in at least two of the lists or databases were kept as the *gene literature list* (259 genes).

### 2.7 Networks & Key driver analysis (KDA)

The networks of different tissues & cell types were downloaded from the GIANT database [56] https://hb.flatironinstitute.org/download. In addition, the precomputed networks *Bayesian_Multitissue* and *PPI* were used directly on the Mergeomics [108] web service (http://mergeomics.research.idre.ucla.edu/samplefiles.php. To remove potential biases associated with network sizes, we only considered the top 1.5 million edges in terms of regulatory scores for each network.

Each of these networks was used to run a key driver analysis (KDA) associated with a list of RA-associated genes. We performed KDA with the Mergeomics R library [57] with a search depth and edge weight set to 1 and 0.5, respectively. In brief, each node (gene) in the network was tested independently. Mergeomics computed the number of edges connecting the node to any gene listed as RA-associated. A node was designated as a key driver if its linkage count exceeded the average number of links to the RA list by more than one standard deviation. In practice, Mergeomics adjusts this number accounting for the regulatory weight associated with each of these links.

### 2.8 Gene regulatory networks in synovial tissues and cell types

We inferred GRNs with PANDA [44]. PANDA combines gene expression profiles of synovial tissues (and cell types) with prior knowledge about TF binding motifs (a list of target genes for each TF) and TF-TF interactions [109, 86]. TF-TF interactions and TF motifs were inferred from the StringDB [43] and CIS-BP database [42], respectively. They were downloaded directly from the GRAND database [110] (https://grand.networkmedicine.org/).

PANDA employs message passing to merge a prior network (derived from mapping TF motifs onto the genome) with protein-protein interaction and gene expression datasets, iteratively refining edge weights in the networks. Applied to our data, PANDA produced directed networks of TFs to their target genes (TGs), comprising 644 TFs and 18 992 genes, resulting in 12 230 848 edges. Here, each edge between a TF and its TG is associated with a weight, which represents the probability of a regulatory interaction between the TF and the TG. The weight values, after undergoing a Z-transformation, typically range between −4 and 4. These indicate the number of standard deviations below (for negative Z-scores) or above (for positive Z-scores) the average weight of the network.

Then, we used LIONESS [45] to estimate an individual gene regulatory network for each individual sample in our RNA-Seq data (see Table S2 for the sample count per cell type). LIONESS estimates sample-specific networks by sequentially leaving each sample out, calculating a network (with PANDA) with and without that sample, and using linear interpolation to estimate the network for the left-out sample. All networks were inferred with the python library netZooPy (https://github.com/netZoo/netZooPy).

### 2.9 Analysis of TFs RA regulatory activity in gene regulatory networks

We leveraged this collection of networks to test whether the weights of these regulatory edges differ significantly between RA and control samples, and to identify the TFs that may potentially be driving these regulatory differences. A Student’s *t*-test was used to estimate: (i) the differential gene expression between RA and the control group, denoted as *t*_expr_; and (ii) the differential weight of the regulatory edges between RA and the control group, denoted as *t*_edge_. We define RA differentially expressed genes as the ones having a |*t*_expr_|> 1.

Note that the *t*-score represents the difference between the mean values of the two groups being compared, divided by the standard error of the difference. A positive (respectively negative) score indicates situations where the mean of the RA group is larger (respectively smaller) than the mean of the control group. The larger the absolute value of the *t*-score, the more statistically significant the difference is relative to the variability of the data. Hence, we quantified the regulatory importance of TFs as the average of the absolute values of the differential weights of the regulatory edges, *t*_edge_ (Eq. 3). We only considered the RA-associated TGs listed in the prior knowledge about TF regulon (CIS-BP database [42]) Defining *G* as the network’s gene set, the above definition can be formalized as follows:

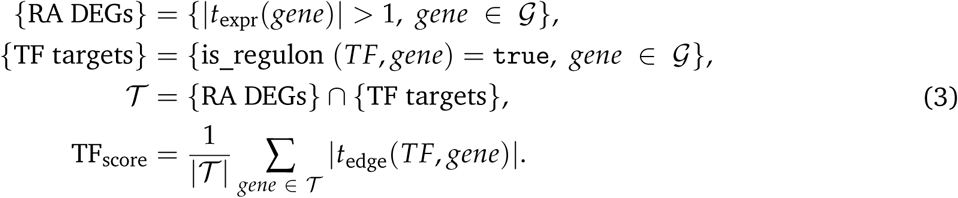

We expect that TFs with the highest scores are the most likely to contribute to RA regulation.

### 2.10 TF-TF co-regulation network

We quantified the co-regulation between TFs by evaluating the correlation of their common TGs’ differential edge weights. TFs with less than 10 common TGs were associated with a co-regulation of 0. Let G be the set of all genes in the network, then:

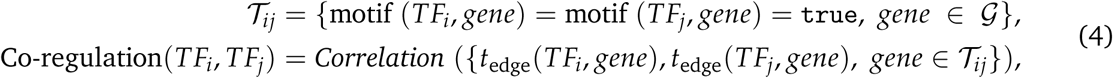

### 2.11 Clustering and Diversity analysis

First, we computed for each cell type a TF-TF distance matrix, defined as one minus the absolute value of the pairwise correlation matrix. Then, in each TF-TF co-regulation network, we clustered TFs into co-regulation groups with a hierarchical agglomerative clustering (HAC) [111]. The clustering criterion was defined as the Ward’s minimum variance method [112], with a distance threshold of 0.5. Ward’s method minimizes the total within-cluster variance, which means at every step, the algorithm finds the pair of clusters that leads to a minimum increase in total within-cluster variance after merging, until the intra-cluster distance is above the threshold.

Then, we characterised these clusters with diversity metrics such as richness (the number of clusters), Shannon entropy and dominance [82]. Dominance is defined as the fractional abundance of the most abundant cluster, while the Shannon entropy (*H*) provides a measure of the overall diversity within the system by considering the frequencies of all clusters, each weighted by the logarithm of its frequency. Denoting *S* the number of clusters, and *p_i_* the fraction of population of cluster *i*, *H* is defined as [113]

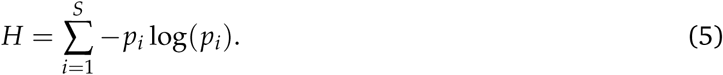

## Supplementary Materials

Supplementary Data 1 and Supplementary Data 2 are available for download.

## Data availability

Gene expression dataset used in this study were taken from GSE89408 [26] and ImmPort (https://www.immport.org/shared/study/SDY998, study accession code SDY998) [15]. All the gene lists obtained in this study, along with the data and the code to reproduce all figures presented in this article are made available publicly on Github at https://github.com/AI-SysBio/RA-drug-discovery. PANDA and LIONESS networks generated in this study will be uploaded upon acceptance.

## Acknowledgments and Funding

The authors thank Camila Lopes-Ramos and Ki-Jo Kim for their valuable suggestions. The work presented in this research received funding from the European Union’s Horizon 2020 research and innovation program under two grant agreements: the COSMIC European Training Network (grant No 765158) and the EU project iPC (grant No 826121). Dr. Gulko and Laragione were funded by the NIH R01AR07316, by the Icahn School of Medicine at Mount Sinai, and by the Eunice Bernhard fund.

## Author contributions statement

A.P. performed all the analysis, under the supervision of M.R.M and P.G. All authors participated in the regular discussions and reviewed the manuscript.

## Supplementary Materials

### A Cellular composition of synovial tissues

#### A.1 Clustering Synovial tissues according to cell-type enrichment scores

We collected a bulk RNA-Seq gene study of synovial biopsies (GSE89408) [26, 105], containing the gene expression profiles of 28 healthy, 152 RA and 22 osteoarthritis (OA) patients over ~25k genes. We selected 13 cells that were previously reported in synovial tissues [15, 16, 14] and leveraged xCell [36] to infer cell-type enrichment scores from gene expression profile of each tissue in the data (Figure S1A). Then, we clustered the tissues in two clusters with hierarchical clustering, by defining the distance between patients and the Euclidean distance between their normalized xCell enrichment scores (Figure S1B).

**Figure S1:**
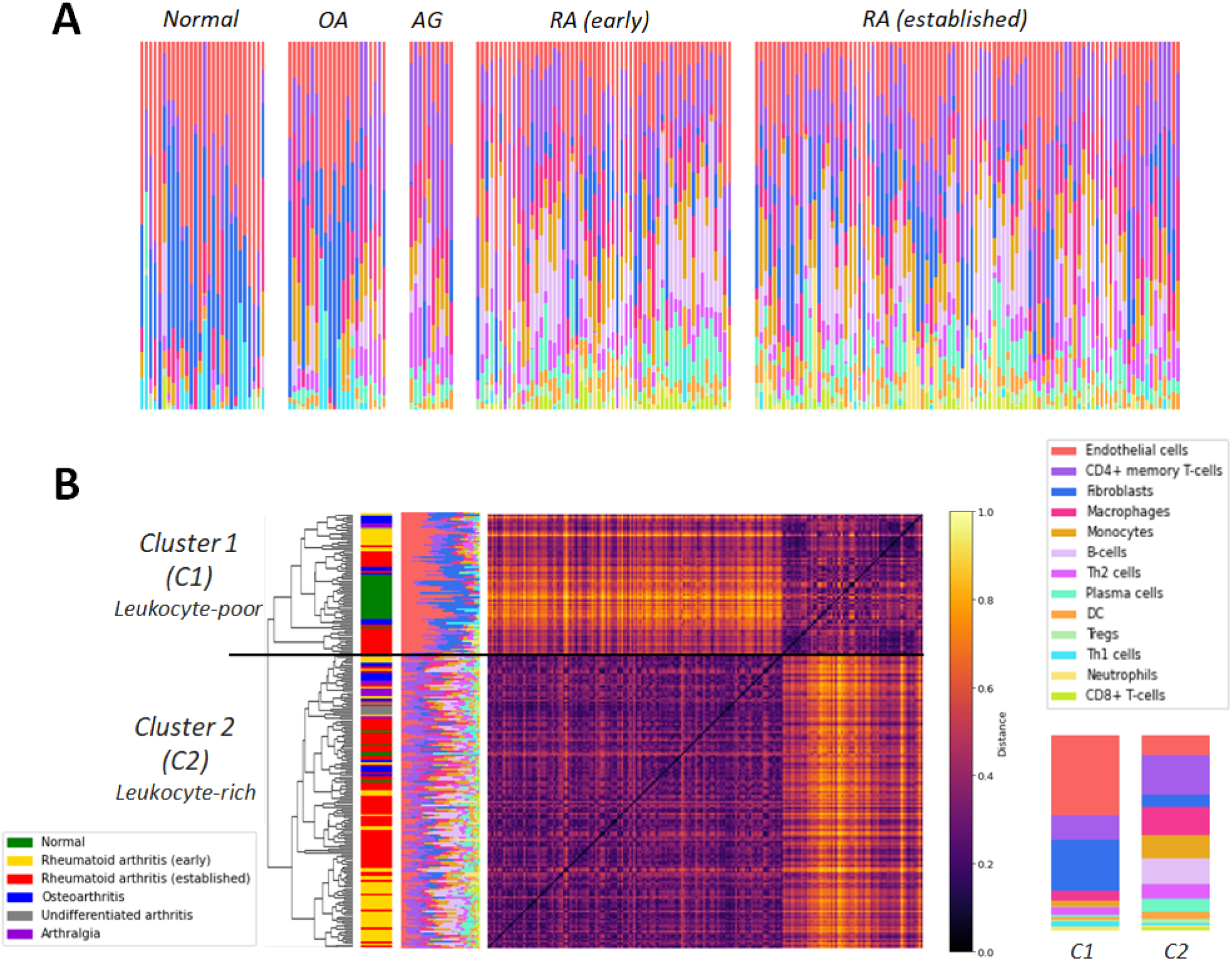
Cellular composition of synovial tissues. (A) The enrichment scores of 13 selected cell types are normalized and displayed in a bar plot for each sample. The samples are grouped according to their pathology: Normal, Osteoarthritis (OA), Arthralgia (AG), early RA and late RA. (B) Heatmap of the normalized Euclidean distance matrix between all synovial tissues, ordered according to their clusters from hierarchical clustering. The mean cellular composition of the two clusters (C1 and C2) is depicted on the right. Note that here we show only 13 cell types for visualization purpose.

We obtain one cluster (C1), mainly characterized by endothelial cells and fibroblasts, while the other one (C2) contains a collection of immune related cell-types. We thus refer to these clusters as Leukocyte-poor and Leukocyte-rich, respectively. While all healthy tissues were clustered in the Leukocyte-poor cluster, a subset of early and established RA tissues were also classified in C1. Zhang & al. [15] observed a similar pattern in their study and they discuss this observation as a potential source of heterogeneity in the effectivity of RA treatments. Interestingly, they find a positive correlation between the number of leukocyte and the level of inflammation in the tissue.

#### A.2 Deconvolution with CIBERSORT

In the main text, we tested for differential enrichment scores across RA and control synovial tissues with xCell [36], which provided an *enrichment score* for each tested cell-types. As deconvolution methods for cellular composition are typically prone to error [114], we verified if our results were consistent with other existing methods. An extensively used pipeline for this task is CIBERSORT [40], which deconvolute directly the cellular composition of the tissues from a *signature matrix* comprised of barcode genes that are enriched in each cell-type of interest (Figure S2). To run CIBERSORT, We used the web-tool CIBERSORx (https://cibersortx.stanford.edu/runcibersortx.php) with a pre-loaded signature matrix comprising 22 immune cells.

**Figure S2:**
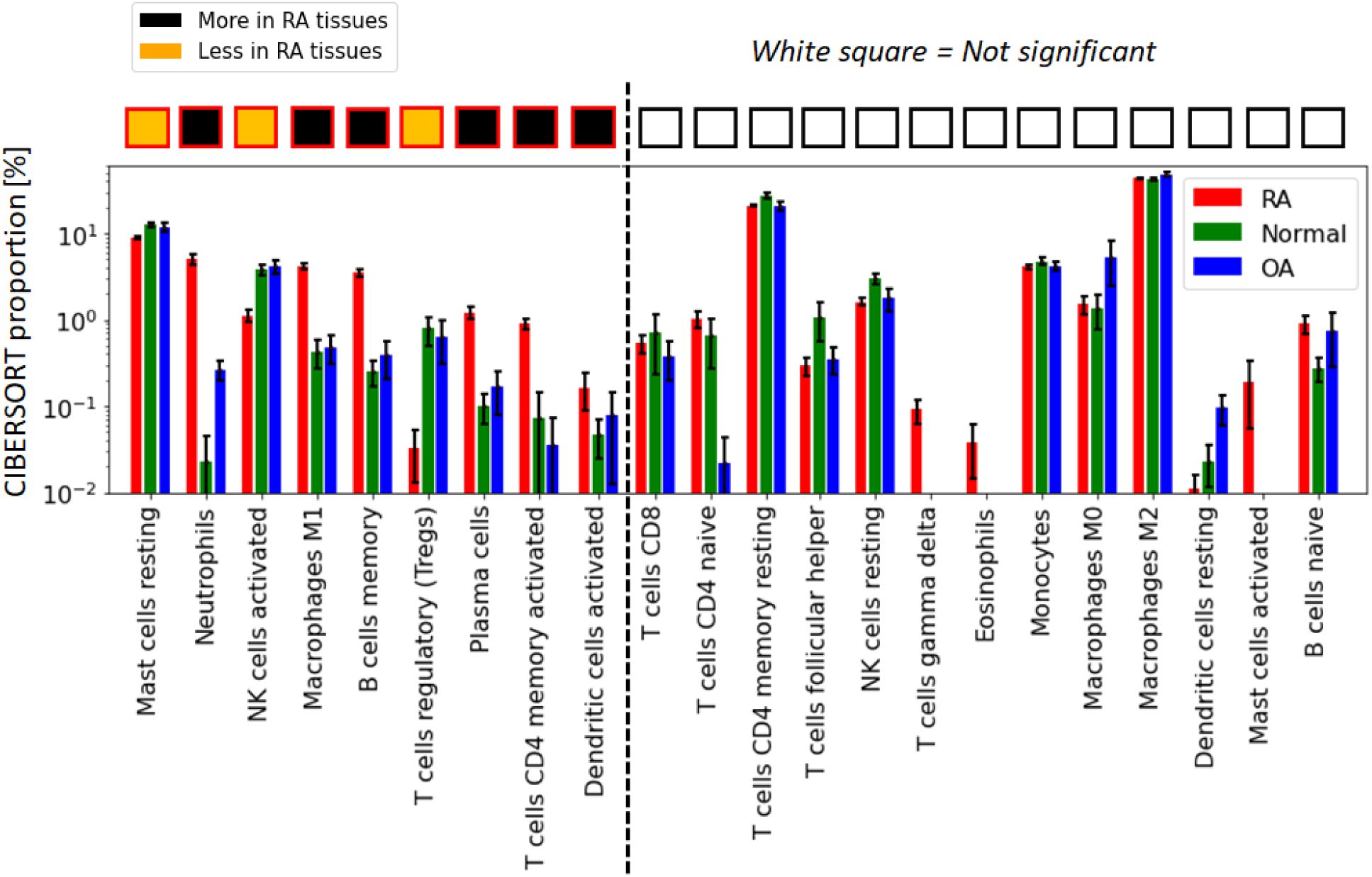
CIBERSORT inferred cellular composition in synovial tissues. cell-type with significantly different composition (*p <* 0.05, *t*-test) in both RAvs normal and RA vs OA tissues are shown on the left. Error bars are 95% confidence intervals defined as 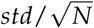.

Overall, we find a good agreement between CIBERSORT and xCell results (Table **??**). The output agrees on Plasma cells, memory B cells, CD4+ memory T cells, and Dentritic cells. On the other hand, a few cells with significant differential enrichment with xCell were not in CIBERSORT (Eosinophils, Monocytes). Note that we could not compare other cell-types in the xCell tool, as the singular matrix for non-immune related cell-types were not available in the CIBERSORTx webtool. A complete comparison would require the construction of a signature matrix comprising all cell-types.

**Table S1:** Agreement between xCell and CIBERSORT cellular deconvolution on relevant cell types. Here we only show cell types that (i) could be evaluated by both tools and (2) that were overrepresented (UP) or underrepresented (DOWN) in RA synovial tissues against both OA and healthy synovial tissues with xCell enrichment scores. NS indicates non significant.

| Cell Type | xCell | CIBERSORT |
| --- | --- | --- |
| Memory B cells | UP | UP |
| Plasma cells | UP | UP |
| CD4+ memory T cells | UP | UP |
| Dendritic cells | UP | UP |
| Monocytes | UP | NS |
| Eosinophils | DOWN | NS |

#### A.3 Explained variance of the RNA expression

A linear regressor (with coefficients denoted as *β*) was fitted to predict the gene expression values from the normalized enrichment scores from xCell (*c_i_*, *i cell-type*), and the explained variance was then quantified as the R-Squared (*R2*) score between the predicted and original gene expression (*Y^predict^* and *Y*, respectively).

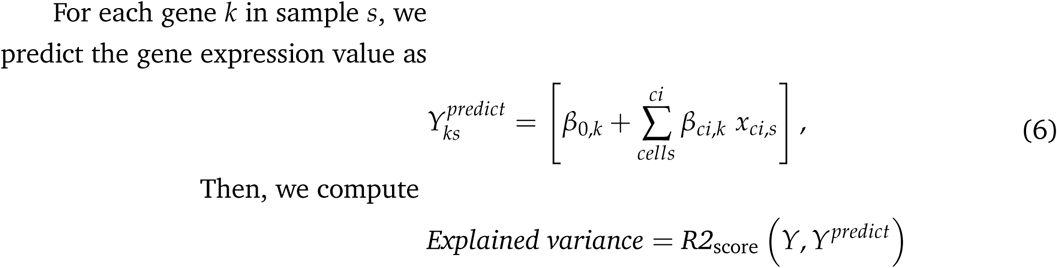

Strikingly, we found that the 18 cell-types discussed in the main text (Figure 1B) cumulatively account for the majority of the variance (*R2* = 61%) in the gene expression data (Figure S3). As expected, the explained variance increases as the enrichment scores of more cell-types are used to predict the gene expression values (from 16% with MSC only to 61% with the 18 cell-types).

**Figure S3:**
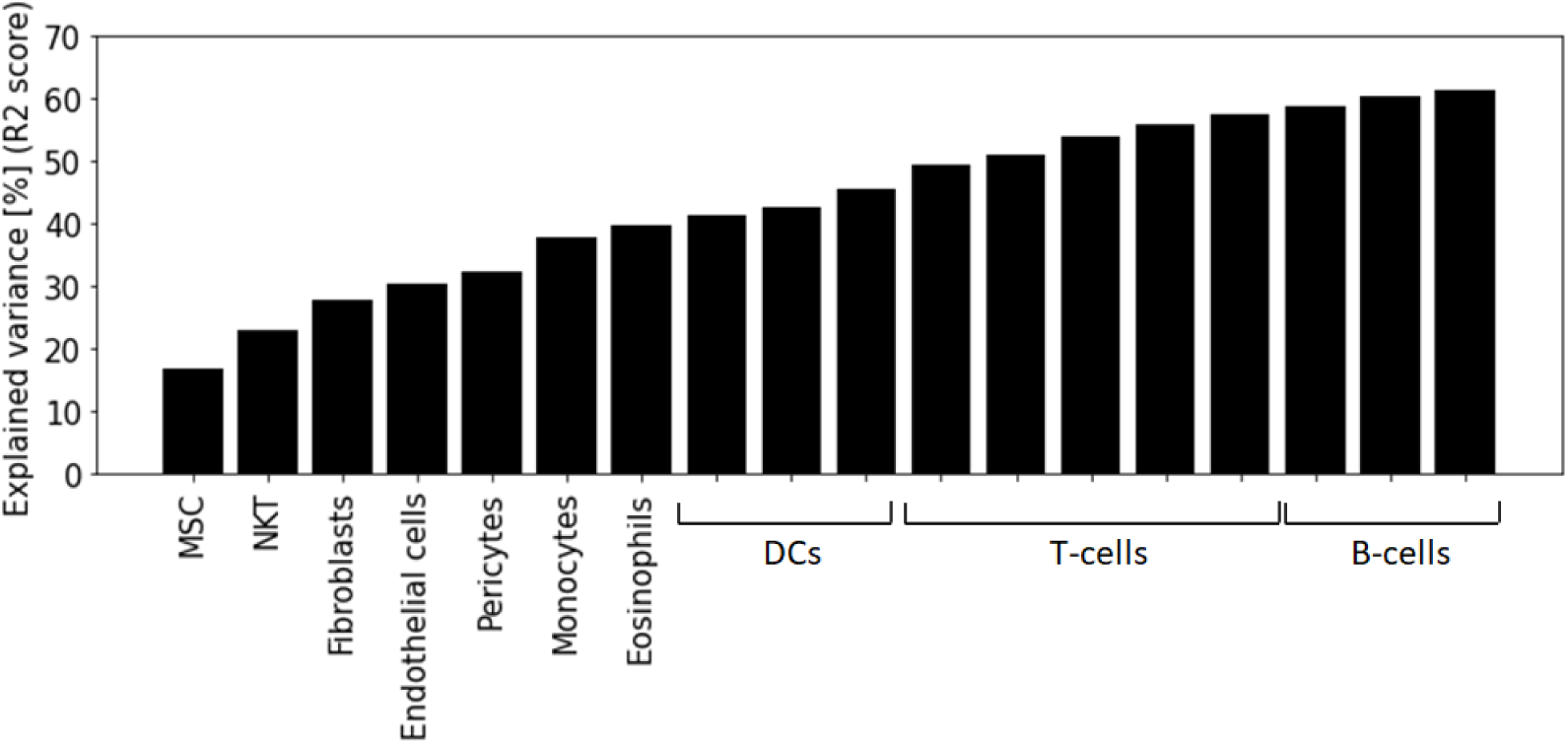
Cumulative explained variance of the cellular composition of the synovial tissue to gene expression. A linear regressor is fitted to predict the gene expression from each gene, where the enrichment score of the cell written in abscise is added to the previous ones (on its left) to train the linear regression.

### B RA associated genes

#### B.1 Differentially expressed genes in synovial tissues

As discussed in the article, the variation of cellular composition in synovial tissue accounted for a large portion of gene expression variability. To capture gene expression variation caused by a changes in actual molecular state rather than changes in cellular composition across samples, we corrected the gene expression profiles to account for the tissue’s cellular composition. More specifically, we considered as confounding factors the enrichment scores of cell types differentially enriched (*p <* 0.05, Student’s *t*-test) between both RA *vs* OA and RA *vs* normal group (18 cell types), which we corrected with a simple linear regression model (Method 2.2 of the main text). Then, we used these corrected gene expression data to perform differential gene expression analysis. Differentially expressed genes (DEGs) were defined as genes obtaining a *p*-value lower than 0.05 in a Student’s *t*-test between the RA and the control group, after an FDR=0.05 correction with the Benjamini & Hochberg procedure [29]. We obtained 279 DEGs.

**Figure S4:**
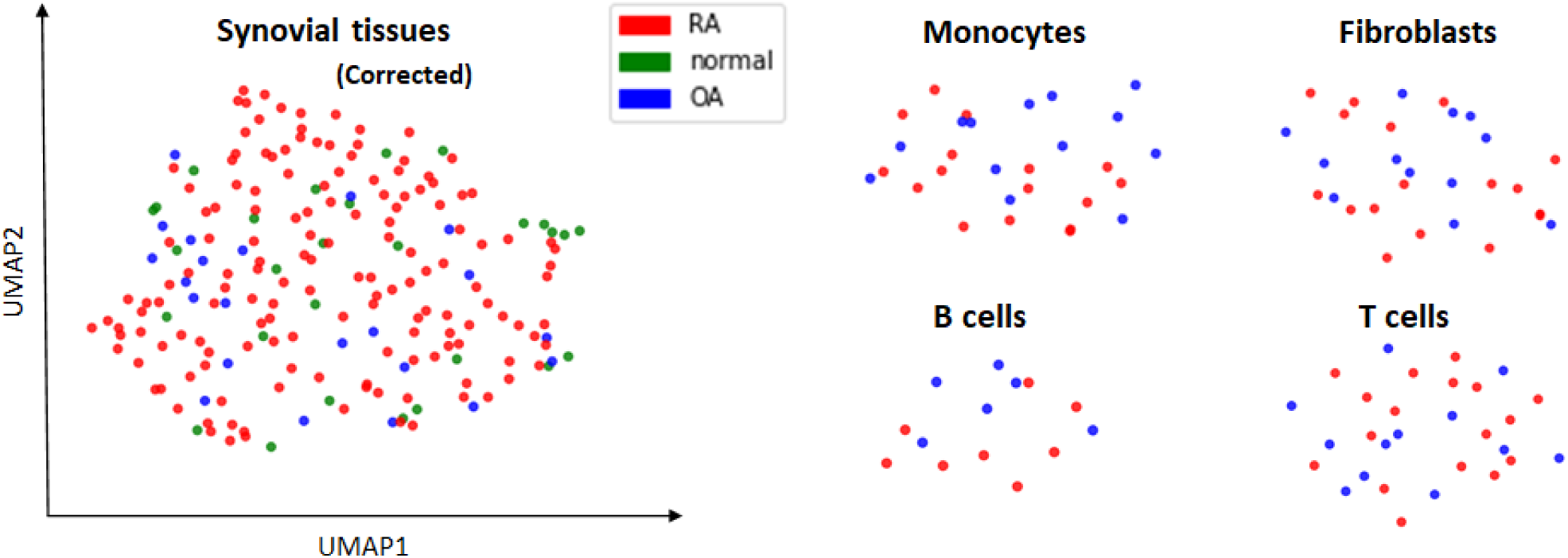
UMAP representation of bulk gene expression data from both cell-type specific and synovial tissues samples. The UMAP representation does not show a clear separation between RA and control groups, indicating a small difference in their gene regulation.

To complement this analysis, we also performed DEG analysis in cell-type specific gene expression data of synovial tissues from [15]. The data consists of both bulk and single-cell gene expression data from monocytes, fibroblasts, B cells, and T cells of RA and OA synovial tissues. As the single-cell data included only 3 control patients (OA), we performed our differential expression analysis on the bulk data. Interestingly, we actually did not find any DEG for these cell types, mainly because only at most ~30 samples were available for each of them. It is consistent with the gene expression data of RA and control overlapping each other in the UMAP and TSNE space (Figure S4). This observation supports the results of the first section of the main text, and confirm that there is limited gene regulatory difference between RA and control patient when the analysis is not biased by cellular composition.

#### B.2 Building a list of RA associated genes from databases & literature

As the DEG list we obtained in our study relies on both (i) the correct estimation of cellular composition in synovial tissues and (ii) the correction for it, it alone cannot provide a reliable list of DEGs. Thus, We combined our DEGs to several meta-analysis [58, 59, 60, 12, 13] from synovial tissues in order to obtain a robust list of DEGs. Interestingly, there were several genes differentially expressed in at least two studies. We kept the 93 genes that were found in at least 2 studies (Figure S5A).

**Figure S5:**
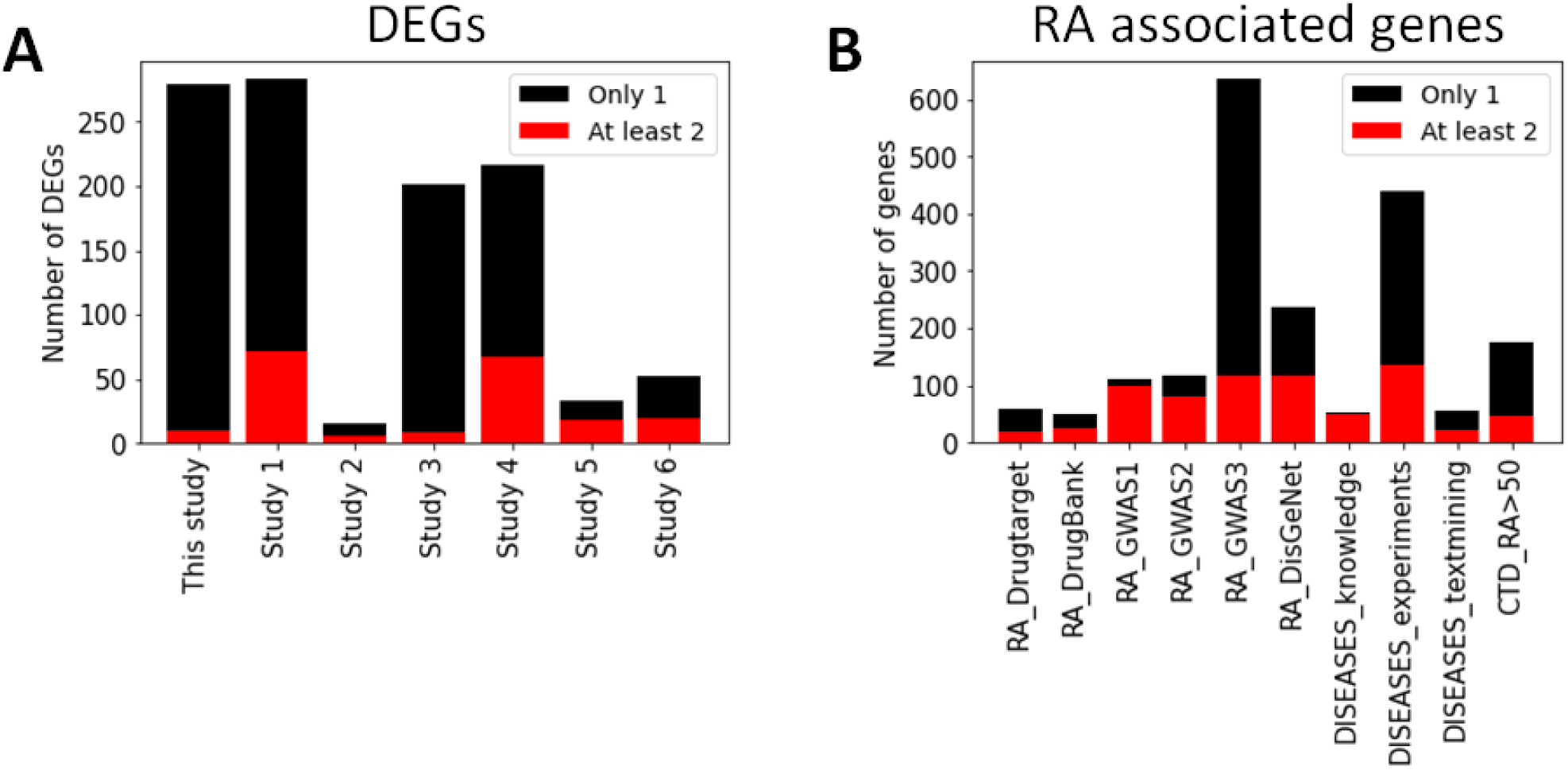
Number of genes across various studies & datasets involved in RA. They are colored depending on if the gene is found only in one study (black) or in at least 2 studies (red). (A) DEGs from various meta-analyses of RA synovial tissues. In addition to our own study, the list of DEG were obtained from Study1 [59], Study2 [60], Study3 [58], Study4 [12], Study5 [13] and Study6 [13] (which contains genes selected with a random forest classifier). (B) RA-associated genes from different databases. Drug_target [66], Drug_Bank [67], GWAS1 [61], GWAS2 [10], GWAS3 [62] (SNP-Disease Associations), DisGeNet [63], DISEASES_knowledge [64], DISEASES_experiments [64], DISEASES_textmining [64] and CTD [65] (kept genes with a score > 50).

Likewise, we performed an extensive literature review to aggregate known genes associated with RA from different contexts. Most recent GWAS revealing genetic risk factors associated with RA were collected from studies [61, 10] and the GWASdb SNP-Disease Associations [62] database. Various genes associated with RA susceptibility were fetched from the publicly available database DISEASES [64], DisGeNet [63] and the Comparative Toxicogenomics Database (CTD) [65] databases (score > 50). We also collected drug targets either already on the market or undergoing clinical trial from review [66] as well as from the DrugBank database [67] (https://go.drugbank.com/categories/DBCAT003604). Genes that were identified in at least two of the lists or databases were kept as the *gene literature list*, containing 259 genes (Figure S5B).

Below we provide the *Literature list* and *DEG list* that we used in our KDA analysis. Genes identified in GWAS studies [61, 10, 62] are shown in bold.

**DEG_list (93 genes with 13 in GWAS):**

RFX5, IRF9, TFEC, DDX60, TNFAIP6, TNFSF10, DRAM1, NMI, TNFAIP8, PLEKHO1, GZMA, IL2RG, SAMSN1, SEL1L3, LEPROTL11, ECE1, TRPV2, ENTPD1, JAK2, SLAMF8, CCL18, LY96, GZMB, TRAF3IP3, AKAP1, LPXN, PTTG1, SYK, LRMP, ANK3, CTSH, S100A8, SDC1, CECR1, AIM2, TAPBPL, GPR65, PRDX6, NKG7, COMMD8, OPN3, NAT1, NAGA, OSBPL3, CD38, IL32, DDX24, TLR7, EVI2A, GUCY1B3, MGAT4A, SEMA4D, CD8A, GALNT6, LPIN3, SLC38A6, PLCG2, QPCT, TXNDC9, IFIT1, RFTN1, TRIM21, TNS3, SRRM2, AREL1, KIAA0125, LRRC15, LCK, DNAJC15, SIRPG, FAS, GZMK, GZMH, SYNGR2, ADAMDEC1, C1GALT1C1, UCP2, FKBP11, IL21R, CXCL9, DENND1B, **RASGRP1, PSMB8, CSF1R, SMCO4, TPD52, HLA-DOB, PRKCB, ALOX5, CLIC1, MYC, IRF8, PTPRC, GSN**.

**Lit_list (259 genes with 189 in GWAS):**

VEGFA, SAA1, CLEC16A, IL1RN, CXCR4, IL19, IL17A, IL4, IL18, CAT, FCGRT, CCL2, HSPD1, LTA, IFNG, MECP2, CXCL10, CD14, CCL5, PIP4K2C, ALB, JAK3, TLR4, IL6, IL10, TP53, TGFB1, SPP1, IL1A, IL1B, CD19, GPX3, CYP3A4, GGT6, CCR5, NFKBIA, CCR7, CXCR3, HMOX1, IL6ST, CASP8, TAP1, CD80, IGF1, RBPJ, CCL20, PADI1, ICAM1, FAS, FLNB, PADI6, CXCL16, AGBL2, COMP, PADI3, IL15, DHODH, IL13, MMP9, SAG, VIM, **CXCR5, TNFRSF1B, EOMES, LPP, MTF1, PRDM1, ILF3, HLA-DRB1, CDK2, IQGAP1, C2, C5, TRAF1, MBL2, BCL2L15, UBASH3A, PXT1, IKZF3, CSF2, IRF8, ARAP1, MSH5, CTLA4, CCR6, C5orf30, IL11RA, TRAF6, LBH, ETS1, RASGRP1, NFKBIL1, ANKRD55, AIRE, KIAA1109, LST1, SMG7, VTCN1, CD40, ABCB1, GSDMB, HLA-DQA1, IFNGR2, CARD8, CXCL13, NOTCH4, PRRC2A, PLD4, FADS2, PTPN11, IRF5, LY6G5C, RCAN1, HLA-DPB1, CFB, HLA-DMB, LOC100506023, CCL19, FOXO1, PRMT1, IL2RB, BACH2, IRF4, CD2, CCL27, PSMB9, PTPN2, NFKBIE, CCL21, PADI4, CLIC1, LOC145837, PRKCQ, BLK, BCL3, DCLRE1C, HLA-DMA, RUNX1, HLA-DPA1, PUS10, FAM107A, BAD, SWAP70, RCOR1, NGF, BAG6, CLNK, APOM, RAD51B, TNFRSF14, UBE2L3, TAGAP, PSMB8, COG6, BAX, ZNF438, TNPO3, CALCR, ATM, GATSL3, AFF3, RELA, CD28, DDX6, IL21, IL20RA, HSPA14, FADS1, RTKN2, TNFAIP3, CD5, TXNDC11, ETV7, PAPOLG, FGFR1OP, FCGR2B, ICOS, STAT4, P2RY10, PANK4, FCGR2A, TEC, REL, HLA-C, ATG5, CDK5RAP2, CFLAR, C1QBP, PDE2A, PRKCB, PTPRC, CD83, PLCL2, ISG20, MICA, SPRED2, TYK2, TAPBP, CEP57, FADS3, ANXA3, FNDC1, IL2RA, PTPN22, IL6R, CD226, TAP2, JAZF1, TPD52, HLA-DOB, IL23R, KIF5A, ARID5B, MYC, CSF3, GATA3, TNF, SYNGR1, ANXA6, FCRL3, DPP4, AHNAK2, CSNK2B, ICOSLG, DOK6, CELF2, AIF1, LY6G6F, ACOXL, NCF2, PADI2, WDFY4, GPANK1, YDJC, MMEL1, CDK6, PPIL4, PXK, IL2, LY6G6C, SLAMF6, CD84, ZMIZ1, PRKCH, FAM167A, SH2B3, HLA-B, IRAK1, HLA-DQB1, HLA-A**.

### C Shared Key driver genes (KDGs) across networks

In the main text, we identified many key driver genes (KDGs) across the tested networks with key driver analysis (KDA). We checked if these KDGs tend to be the same across networks or if on the other hand, the identified genes across networks were independent. Our results, displayed in Figure S6, indicated that a subset of KDGs were found consistently in the majority of the network. More precisely, the top 20 genes were found in 75% of the networks and more than 500 were found in more than half of the tested networks. This is much higher than what we would expect if the KDG were independent (red line in Figure S6).

**Figure S6:**
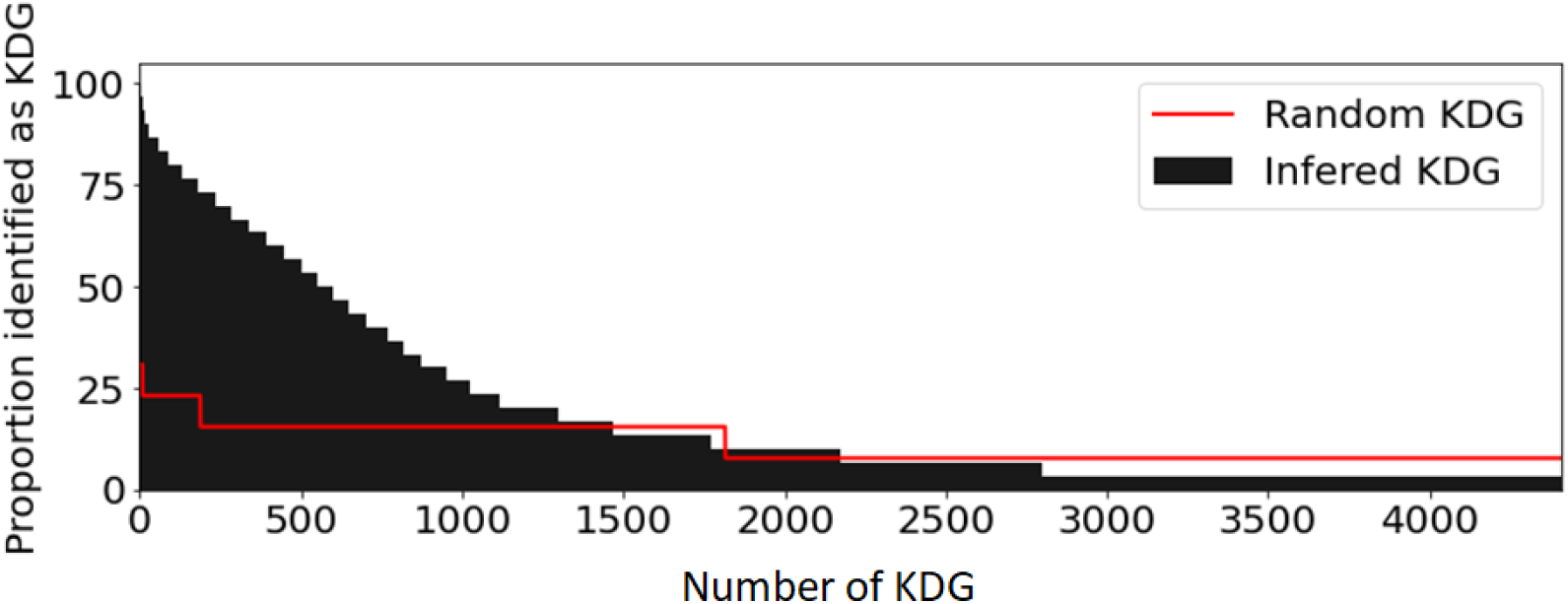
Gene distribution of the proportion of networks in which a given gene was found. The most commons KDGs are found in 80% of the networks while only 25% would be expected if genes were randomly assigned as KDG (red line).

**Top 100 KDG list** (ordered from highest to lowest score, 11 KDGs were also in GWAS [61, 10, 62] marked in bold): PTPN6, HLA-E, HLA-F, GBP1, LCP2, GLIPR1, **HLA-A**, CTSS, SRGN, CTSH, CASP1, **HLA-B**, **HLA-C**, CD44, IFI30, TNFAIP8, CCL5, **PSMB8**, TAP1, ICAM1, ARHGDIB, PSME1, B2M, THEMIS2, SP100, MYD88, **TNFAIP3**, PLAUR, ARPC1B, ITGB2, **HLA-DMA**, CYBA, HLA-G, **PSMB9**, CXCL10, IL7R, CTSC, BCL2A1, CTSB, TNFSF10, SP110, CD86, CD48, CORO1A, PSMB10, PLEKHO2, LYN, EVI2B, TPP1, IFITM2, **ISG20**, NCF4, BTN3A3, IFITM3, UBE2L6, **CFLAR**, MAN2B1, STK10, **NCF2**, WARS1, CD53, IFITM1, FAS, RGS19, LAPTM5, **PLEK**, RAC2, CD74, TNFAIP2, PECAM1, PLAAT4, **HLA-DQB1**, BIRC3, TIMP1, IL1B, NFKBIA, IRF1, STAT1, PHF11, IFIT2, COTL1, NMI, PLSCR1, OAS1, MX2, IFNGR1, CAPG, LGALS3, **NFKBIE**, HMOX1, RAB27A, RAB31, GBP2, CCL2, FYN, **TNFRSF1B**, ADGRE5, CXCR4, SERPINB1, LITAF.

### D Supplementary Figures and Tables

**Table S2:** List of networks involved in our study. GIANT RIMBANET and PPI were downloaded from a public database while PANDA and LIONESS networks were computed specifically for this study. H refers to healthy donors.

| Network(s) name | Type | Method | Used for | Number of networks |
| --- | --- | --- | --- | --- |
| PANDA_Synovial_Tissue | Bipartite | PANDA, LIONESS* | TF reg. score | 152 RA, 22 OA |
| PANDA_FLS | Bipartite | PANDA, LIONESS* | TF reg. score | 18 RA, 12 OA |
| PANDA_Synovial_Monocyte | Bipartite | PANDA, LIONESS* | TF reg. score | 17 RA, 13 OA |
| PANDA_Synovial_Bcell | Bipartite | PANDA, LIONESS* | TF reg. score | 8 RA, 7 OA |
| PANDA_Synovial_Tcell | Bipartite | PANDA, LIONESS* | TF reg. score | 17 RA, 13 OA |
| GIANT_DC | Bayesian | GIANT<sup>§</sup> | KDA | 1 |
| GIANT_NKT | Bayesian | GIANT<sup>§</sup> | KDA | 1 |
| GIANT_Monocyte | Bayesian | GIANT<sup>§</sup> | KDA | 1 |
| GIANT_Tcell | Bayesian | GIANT<sup>§</sup> | KDA | 1 |
| GIANT_Bcell | Bayesian | GIANT<sup>§</sup> | KDA | 1 |
| GIANT_Fibroblast | Bayesian | GIANT<sup>§</sup> | KDA | 1 |
| GIANT_Adipocyte | Bayesian | GIANT<sup>§</sup> | KDA | 1 |
| GIANT_Tonsil | Bayesian | GIANT<sup>§</sup> | KDA | 1 |
| GIANT_LN | Bayesian | GIANT<sup>§</sup> | KDA | 1 |
| GIANT_Blood | Bayesian | GIANT<sup>§</sup> | KDA | 1 |
| GIANT_Spleen | Bayesian | GIANT<sup>§</sup> | KDA | 1 |
| RIMBANET_Multitissue | Bayesian | RIMBANET<sup>†</sup> | KDA | 1 |
| PPI | Undirected | StringDB<sup>†</sup> | KDA | 1 |

\* Computed in this study from bulk RNAseq [26, 15].
<sup>§</sup> GIANT networks downloaded from https://hb.flatironinstitute.org/download.
<sup>†</sup> RIMBANET and PPI downloaded from http://mergeomics.research.idre.ucla.edu/samplefiles.php.

**Figure S7:**
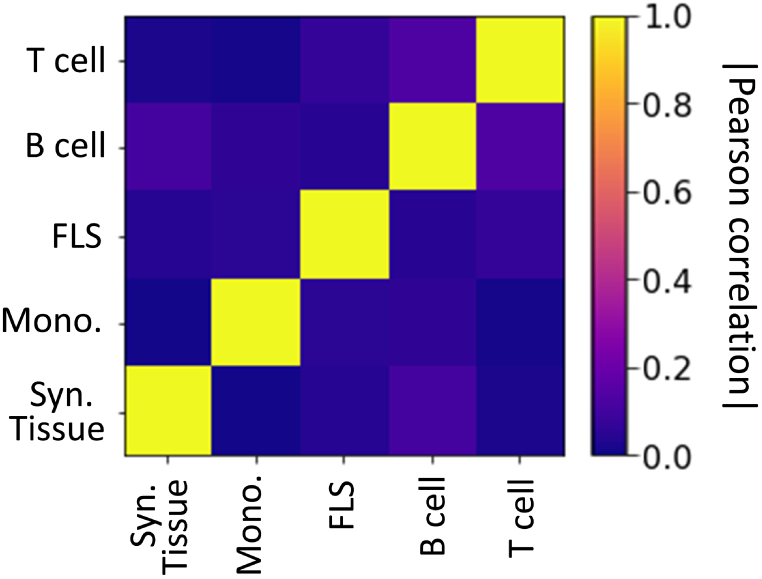
Heatmap of the Pearson correlation between the differential gene expression (t-score) in each tissue type (Synovial tissue, monocyte, FLS, B cell, and T cell.

**Figure S8:**
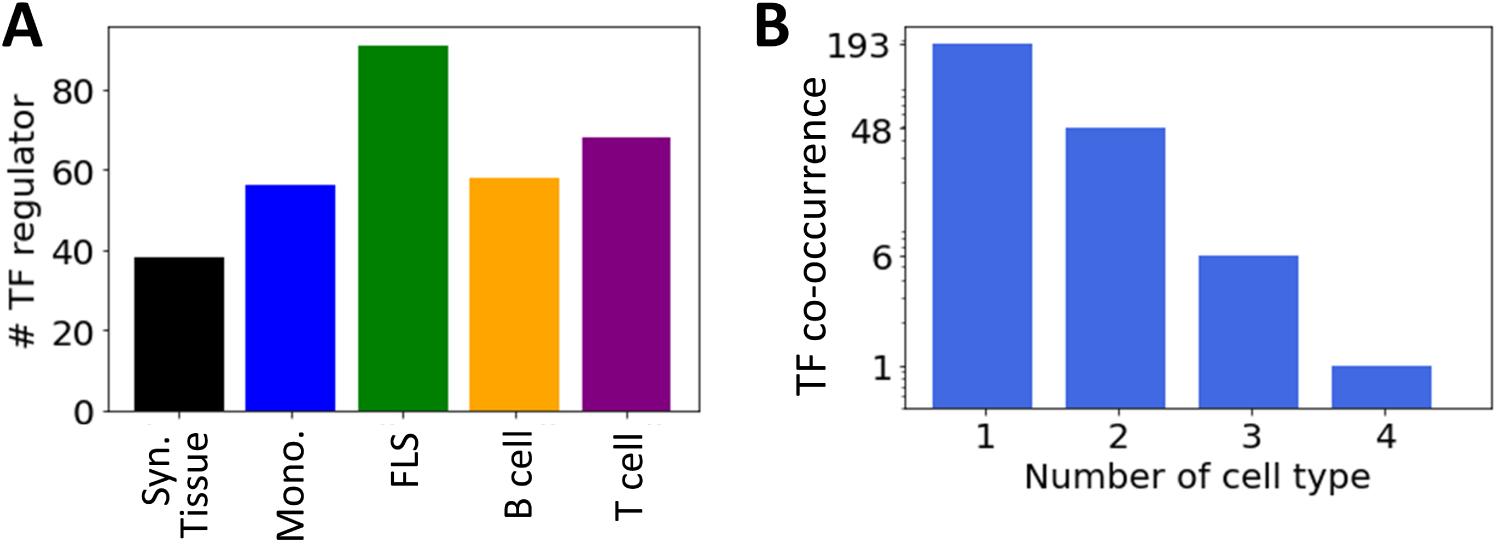
(A) Number of key regulator TFs identified in each cell type. (B) Number of cell type TFs regulators are identified into (note the log-scale on the *y*-axis).

**Figure S9:**
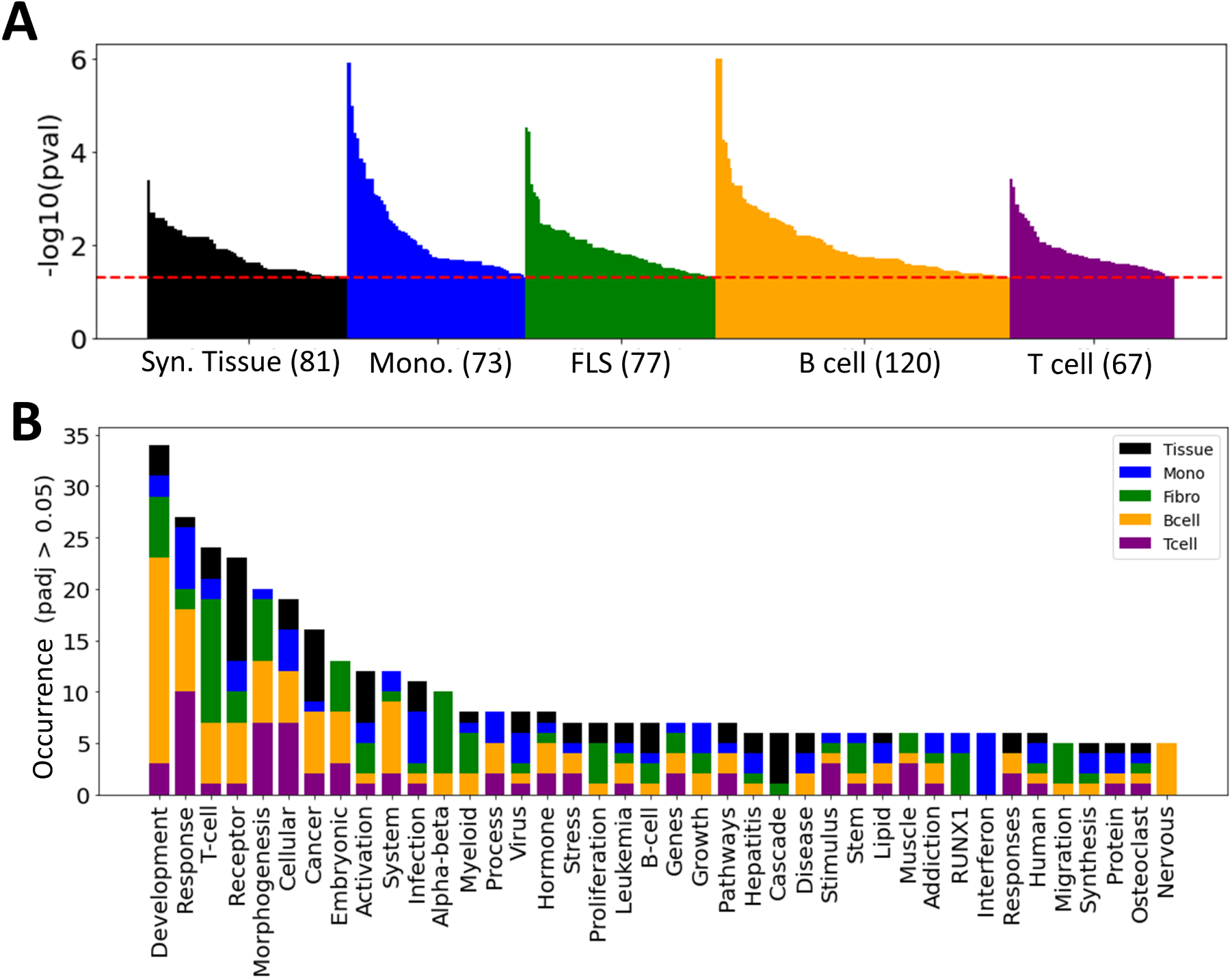
(A) Number of differentiated pathways. (B) Occurence of each word in all the pathway terms combined.

**Figure S10:**
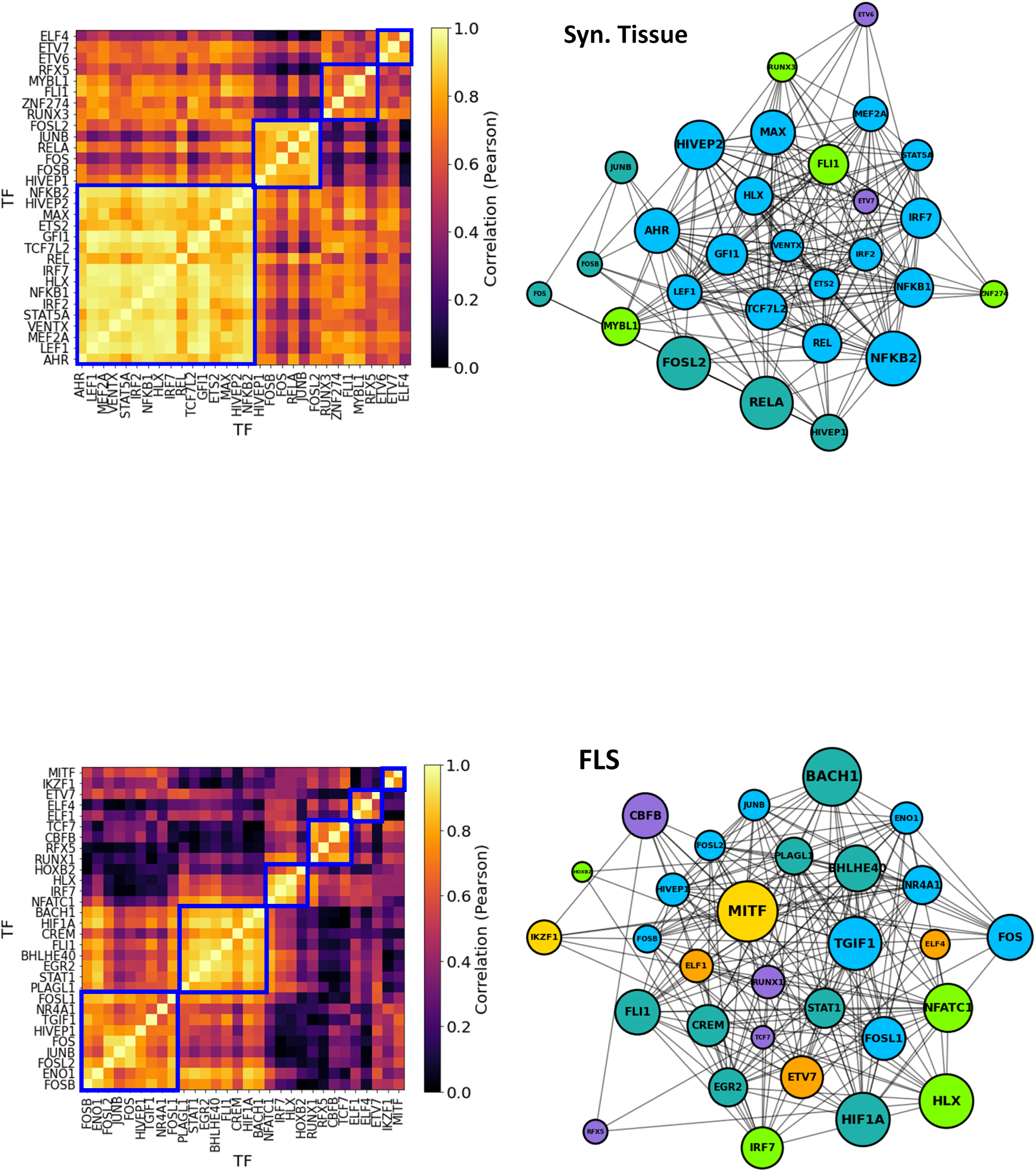
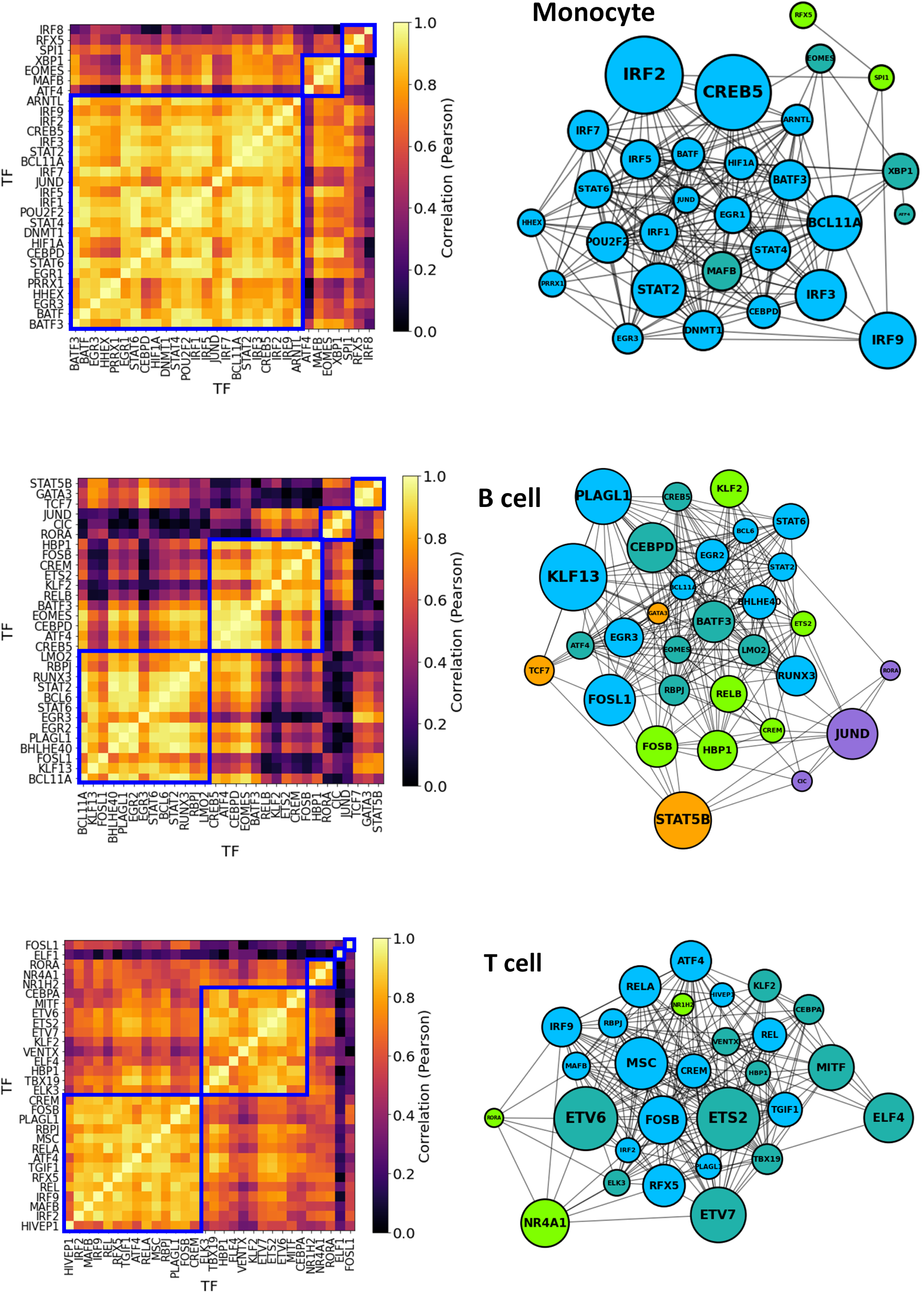
TF-TF co-regulation networks of each cell type analyzed in this article. Network showing the major TFs involved in FLS RA regulation, with nodes size proportional to both the degree and the TF regulatory score (*t*_reg_). Nodes are colored according to their clusters.

